# Anti-inflammatory macrophages limit endothelial inflammation and restrict neutrophil recruitment through Gas6/AXL signaling

**DOI:** 10.64898/2026.10.12.758565

**Authors:** Aitana Ignes-Romeu, Rupashi Sood, Laurel E. Hind

**Affiliations:** Department of Chemical and Biological Engineering, University of Colorado Boulder, Boulder, CO 80303, USA

## Abstract

Neutrophil extravasation requires coordinated interactions between the endothelium and surrounding cells, including tissue resident macrophages. Macrophages adopt distinct polarization states in the tissue environment, positioning them as regulators of vascular inflammation. However, how these states influence barrier function and neutrophil extravasation remains unclear. Here, we polarized macrophages in the matrix surrounding an endothelial vessel, in a three-dimensional inflammation-on-a-chip model developed only from primary human cells. We found anti-inflammatory macrophages limited endothelial inflammation, reducing ICAM-1 upregulation, maintaining VE-Cadherin expression, halving permeability relative to pro-inflammatory devices, and reducing neutrophil extravasation and migration relative to macrophage-free devices. Secretome profiling identified Gas6, an AXL ligand, as a potential regulator of these effects. Inhibiting AXL increased extravasation in devices with anti-inflammatory macrophages, and soluble Gas6 reduced extravasation in macrophage-free devices, implicating Gas6 as a mediator of this regulation. This indicates macrophages regulate endothelial inflammation and the neutrophil response via Gas6 signaling.

**One-sentence summary:** Anti-inflammatory macrophages limit neutrophil entry into tissue through Gas6/AXL signaling and reduce endothelial activation.

## INTRODUCTION

Neutrophil recruitment to sites of sterile injury or infection is a cornerstone of innate immunity, and when this response is dysregulated it drives tissue damage (*1*, *2*). An early and critical step in neutrophil recruitment is extravasation, the process by which neutrophils adhere to the endothelial cells lining a blood vessel and then cross the vessel wall into the surrounding tissue. Extravasation depends on the ordered engagement of endothelial adhesion molecules, primarily intercellular adhesion molecule-1 (ICAM-1) and vascular cell adhesion molecule-1 (VCAM-1), along with remodeling of the intercellular endothelial junctions held together by vascular endothelial cadherin (VE-Cadherin) (*3–6*). The process by which neutrophils adhere on and cross the vessel wall is well understood, yet how tissue-resident macrophages alter the balance of adhesion and junctional proteins, and thereby endothelial barrier inflammation and neutrophil extravasation, remains poorly defined (*7*, *8*).

Blood vessels are surrounded by a stromal layer in which tissue-resident macrophages reside (*9*). In vivo, macrophages positioned alongside and within vessel walls regulate neutrophil traffic: they sense damage-associated molecular patterns released during sterile inflammation, act on endothelial cells to alter vascular permeability, and establish local gradients of adhesion molecules that permit neutrophil transit (*10–15*). Macrophages are also highly plastic, adopting distinct functional states depending on the signals in their environment (*16*, *17*). Macrophages activated by interferon-γ and bacterial products, termed classically activated or pro-inflammatory, express the co-stimulatory molecule CD80 and secrete interleukin-6 (IL-6) and TNF-α, both of which reduce endothelial junction strength (*18*). Macrophages activated by IL-4, termed alternatively activated or anti-inflammatory, express the scavenger receptor CD163 and release pro-resolving mediators such as macrophage-derived chemokine (MDC/CCL22) and soluble CD163 (*19*, *20*). Whether the microenvironment established by either phenotype is, on its own, sufficient to differentially regulate the endothelial barrier, and the magnitude of any such effect, has not been established in a fully human system.

Two obstacles have made this difficult to test directly. The first is that macrophage–endothelial interactions are difficult to isolate in vivo, as inflammatory signaling influences both cell populations at the same time. Furthermore, multiple inflammatory pathways result in the same functional response, making it hard to decipher specific roles (*21*). Mouse models also differ from humans in the neutrophil compartment itself. Neutrophils account for roughly half to three quarters of circulating leukocytes in humans but a far smaller fraction in mice, and mice have no structural homolog of IL-8 (*22*, *23*). We chose IL-8 for this study because of its central role in human neutrophil recruitment, which cannot be studied in mice. The second is that an in vitro model of neutrophil extravasation must reproduce the layered chemotactic fields found in tissues, where neutrophils sense several chemoattractants at once, among them interleukin-8 (IL-8) and leukotriene B_4_ (LTB_4_) (*24*, *25*). IL-8 recruits neutrophils over long distances, whereas LTB_4_ acts as a relay signal that widens the effective recruitment radius and reinforces the front–rear polarity of migrating neutrophils (*26*, *27*). Microphysiological systems (*28*, *29*), especially those built around three-dimensional lumens patterned in extracellular matrix hydrogels (*30*, *31*), allow such an environment to be modeled with defined cell composition and controlled inflammatory timing. These platforms have been used to probe cell–cell crosstalk at the vascular–tissue interface in synovial inflammation (*32*), tumor microenvironment remodeling (*33*), and neutrophil–endothelial extravasation (*34*). Using such systems, our group found that THP-1 macrophages limit neutrophil extravasation in a model bacterial infection (*35*) and that monocytic myeloid-derived suppressor cells impair neutrophil recruitment and migration in the infectious microenvironment (*36*).

Here, we used a three-dimensional inflammation-on-a-chip model to determine how macrophage polarization regulates endothelial activation and the neutrophil response during sterile inflammation. Primary human monocytes were differentiated and polarized in situ within a collagen–fibronectin matrix surrounding an endothelial lumen. The endothelium was then primed with TNF-α to model sterile inflammation, and primary human neutrophils were introduced into the vessel and stimulated to migrate up a combined IL-8 and LTB_4_ gradient. We found that anti-inflammatory macrophages reduced endothelial activation and limited neutrophil extravasation, whereas pro-inflammatory macrophages had the opposite effect on the endothelium. Finally, we identified Gas6 as a negative mediator of neutrophil extravasation that is enriched in the presence of anti-inflammatory macrophages. Collectively, these findings demonstrate that macrophage polarization state plays a regulatory role in endothelial inflammation and neutrophil recruitment.

## RESULTS

### Primary human macrophages polarize and remain viable within the microphysiological device

Before we could determine how macrophage polarization regulates the inflammatory microenvironment, we first needed to integrate primary human macrophages into our device and confirm their viability and polarization. Monocytes were embedded in a collagen–fibronectin hydrogel around a hollow lumen, differentiated with M-CSF for five days, and polarized in situ on Day 6. On Day 7, endothelial cells were added to the hollow lumen to create a model vessel (Fig. 1A). Vessels formed over the following two days, and extravasation experiments were performed on Day 9.

**Fig. 1.**
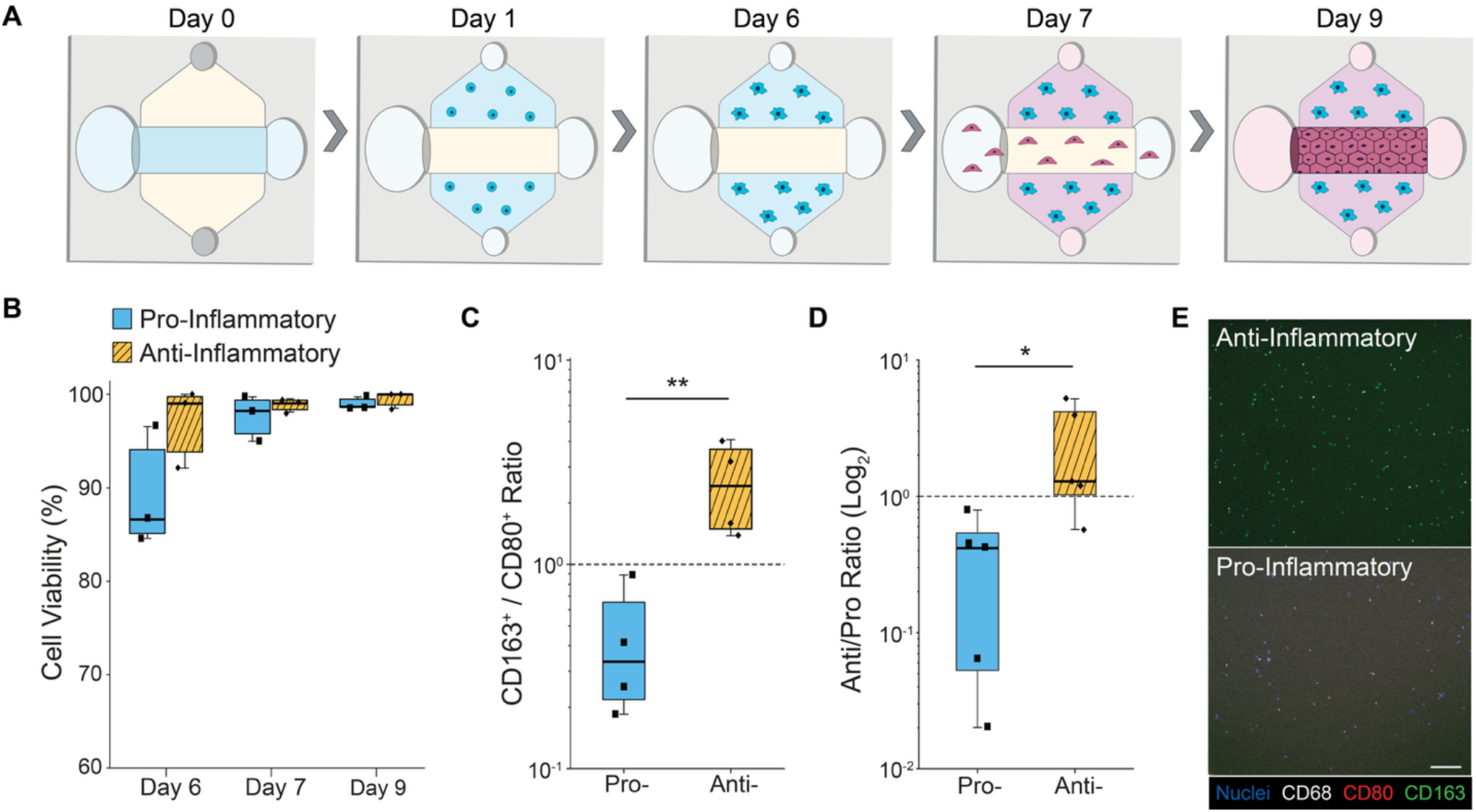
Inflammation-on-a-chip platform and phenotypic validation of primary human macrophage microenvironments. (A) Schematic of the three-dimensional inflammation-on-a-chip device. A cylindrical lumen lined with human umbilical vein endothelial cells sits within a collagen–fibronectin extracellular matrix. Primary human monocyte-derived macrophages are polarized in situ within the matrix toward pro-inflammatory (Pro-) or anti-inflammatory (Anti-) phenotypes. Top and side ports allow injection of chemoattractants, neutrophils and inhibitors. (B) Cell viability (%) for pro- and anti-inflammatory macrophages on Days 6, 7 and 9. N = 3, n = 3. (C) Polarization Index (CD163^+^/CD80^+^ ratio, log scale). Welch’s t-test on log_2_-transformed values, P = 0.007. N = 4–6, n = 3. (D) Anti/Pro cytokine secretion ratio (log_2_) for pro- and anti-inflammatory macrophages in 3D collagen– fibronectin hydrogels, quantified by multiplex MagPix Luminex assay. The ratio is the summed concentrations of the three anti-inflammatory analytes (IL-10, soluble CD163 and MDC/CCL22) divided by the summed concentrations of the two pro-inflammatory analytes (IL-6 and TNF-α). Welch’s t-test on log_2_-transformed values, P = 0.0275. N = 4– 5, n = 1. (E) Representative confocal maximum-intensity projections of anti-inflammatory (top) and pro-inflammatory (bottom) macrophages in the 3D collagen–fibronectin matrix on Day 9, stained for Hoechst (nuclei, blue), CD68 (white), CD80 (red) and CD163 (green). Scale bar, 100 µm. In (B) to (D), the center line is the median, the box spans the interquartile range and the whiskers span the full rang(E) with individual biological replicates overlaid.

Macrophage viability remained high for both pro- and anti-inflammatory phenotypes, with approximately 99% of macrophages viable on Day 9 (Fig. 1B). We next determined whether the embedded monocytes could be polarized into two distinct populations, one biased toward a pro-inflammatory phenotype and one toward an anti-inflammatory phenotype. On Day 9, polarized macrophages were immunostained and imaged by confocal microscopy for CD80 (pro-inflammatory), CD163 (anti-inflammatory) and CD68 (pan-macrophage) (Fig. 1E). CD80^+^ and CD163^+^ cells were quantified as a percentage of CD68^+^ cells, and the two percentages were combined into a Polarization Index (%CD163^+^/%CD80^+^). A value above 1.0 indicates a population biased toward the anti-inflammatory phenotype and below 1.0 one biased toward the pro-inflammatory phenotype. Every anti-inflammatory replicate fell above 1.0 and every pro-inflammatory replicate below it, so the two conditions produced populations with opposing phenotypes (Fig. 1C).

We also confirmed macrophage polarization by secreted protein analysis to quantify differences in the signaling environments generated by the two cell populations. Pro- and anti-inflammatory macrophages were embedded in collagen–fibronectin hydrogels, media was conditioned for 24 h, and the conditioned media was profiled using a multiplex bead-based immunoassay. Five analytes were quantified, IL-6 and TNF-α as pro-inflammatory markers and IL-10, soluble CD163 and MDC/CCL22 as anti-inflammatory markers. For each replicate, we calculated the ratio of the summed concentrations of the three anti-inflammatory analytes to the summed concentrations of the two pro-inflammatory analytes, so that the ratio reports the balance of secreted mediators. Anti-inflammatory macrophages had a median ratio above 1.0 and pro-inflammatory macrophages below 1.0 (Fig. 1D, log_2_ scale; absolute analyte concentrations are reported in fig. S1). Pro- and anti-inflammatory macrophages therefore maintain distinct functional states and generate chemically distinct secretomes within an identical extracellular matrix.

### Macrophage polarization differentially regulates endothelial ICAM-1 and VE-Cadherin expression

To determine how the presence of pro- or anti-inflammatory macrophages alters endothelial adhesion and junctional protein expression, we measured ICAM-1 and VE-Cadherin both with and without inflammatory stimulation. Expression levels were normalized to nuclei count and expressed as log_2_ fold change (log_2_ FC) relative to each replicate’s own unstimulated, macrophage-free condition (Fig. 2A). Stimulation here was TNF-α introduced into the lumen for 2 h. Five conditions were compared in our devices. These were a stimulated macrophage-free control (No Mac.(+)), devices with pro-inflammatory macrophages with or without stimulation (Pro(+), Pro(−)), and devices with anti-inflammatory macrophages with or without stimulation (Anti(+), Anti(−)) (Fig. 2).

**Fig. 2.**
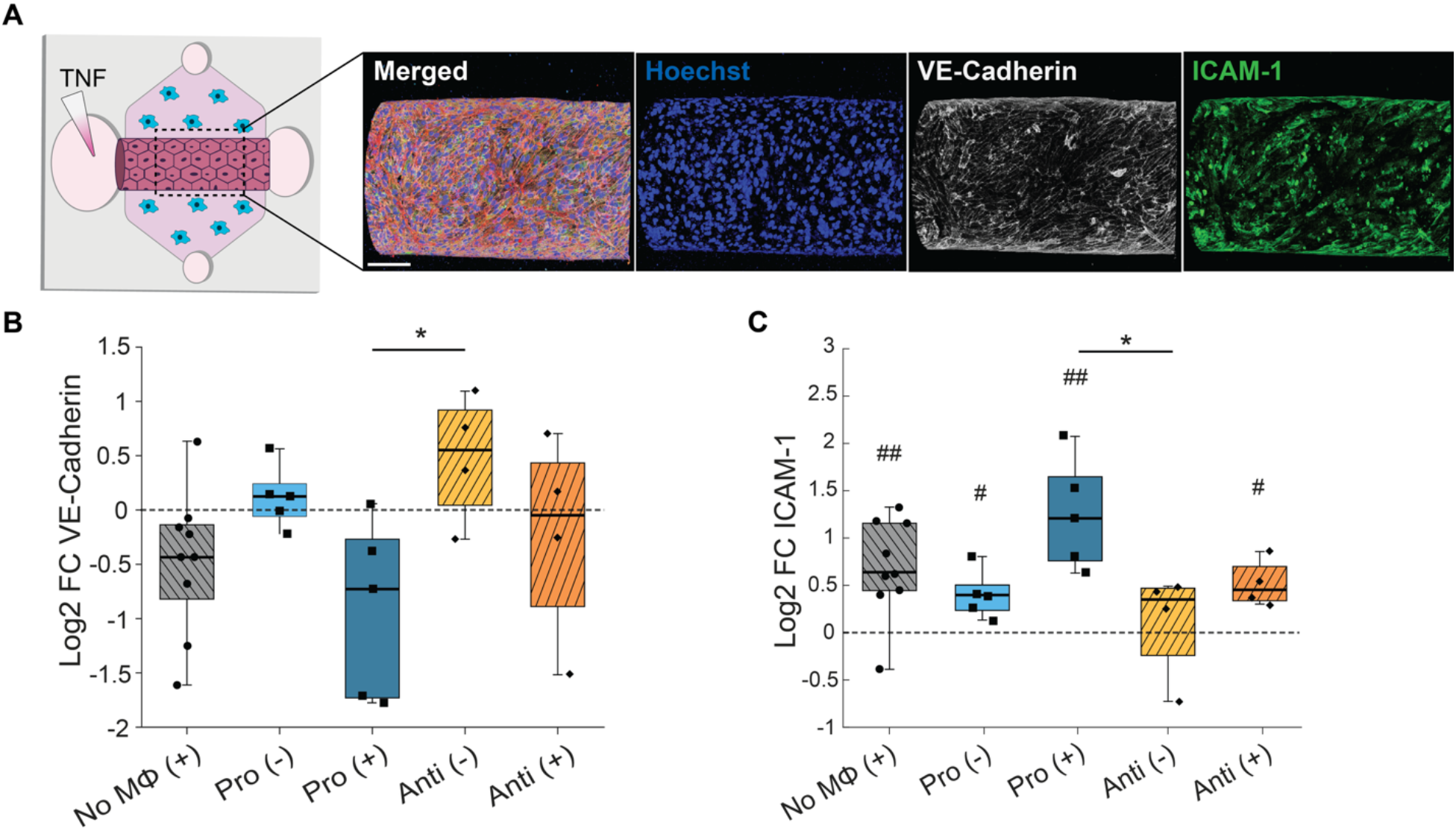
Pro- and anti-inflammatory macrophage microenvironments differentially regulate endothelial ICAM-1 and VE-Cadherin. (A) Schematic of the immunofluorescence workflow. On Day 9 the HUVEC-lined lumen is fixed and stained for VE-Cadherin, ICAM-1, F-actin and nuclei, a custom MATLAB segmentation generates a vessel mask, and total signal within the mask is summed per image, normalized to nuclei count, and expressed as log_2_ FC relative to each replicate’s own unstimulated, macrophage-free condition. Scale bar, 150 µm. (B) Log_2_ FC of VE-Cadherin across the five tested conditions. (C) Log_2_ FC of ICAM-1. One-way ANOVA with Tukey–Kramer post hoc correction for comparisons between conditions, and a one-sample t-test of each condition against baseline (log_2_ FC = 0). Between conditions, *P < 0.05 and **P < 0.01. Against baseline, #P < 0.05 and ##P < 0.01. N = 4–9 per condition (No Mac.(+), N = 9; each pro-inflammatory condition, N = 5; each anti-inflammatory condition, N = 4), n = 2–3. In (B) and (C), the center line is the median, the box spans the interquartile range and the whiskers span the full range, with individual biological replicates overlaid. In this figure, (+) denotes stimulation with TNF-α in the lumen for 2 h, without the chemoattractant source, and (−) its absence.

We first measured VE-Cadherin, the adherens junction protein that maintains the endothelial barrier (Fig. 2B). No condition differed significantly from the unstimulated, macrophage-free condition (Fig. 2B). In macrophage-free vessels, inflammatory stimulation produced a small decrease in VE-Cadherin expression (No Mac.(+), median log_2_ FC ≈ −0.43) consistent with TNF-α regulating endothelial junctions. Vessels with pro-inflammatory macrophages showed the greatest decrease in VE-Cadherin expression following stimulation (Pro(+), median log_2_ FC ≈ −0.73) and remained near baseline without stimulation (Pro(−) ≈ +0.12). Vessels with anti-inflammatory macrophages had the highest VE-Cadherin expression when unstimulated (Anti(−) ≈ +0.55) and remained near baseline when stimulated (Anti(+) ≈ −0.05). The only significant difference between conditions was between stimulated vessels with pro-inflammatory macrophages (Pro(+)) and unstimulated vessels with anti-inflammatory macrophages (Anti(−)) (Fig. 2B).

Expression of ICAM-1, the adhesion molecule that mediates neutrophil adhesion, showed the opposite response (Fig. 2C). ICAM-1 expression was enhanced in four of the five conditions compared to the unstimulated, macrophage-free control (marked # in Fig. 2C). Stimulation of macrophage-free vessels increased ICAM-1 expression (No Mac.(+), median log_2_ FC ≈ +0.64, ##). In the presence of pro-inflammatory macrophages, there was a modest but significant increase in ICAM-1 expression (Pro(−) ≈ +0.40, #), and expression increased further with stimulation (Pro(+) ≈ +1.21, ##). In the presence of anti-inflammatory macrophages, unstimulated vessels had ICAM-1 expression comparable to unstimulated macrophage-free controls, and this was the only condition that did not differ significantly from that control (Anti(−) ≈ +0.35). Following stimulation, ICAM-1 expression in these devices increased to a level comparable to unstimulated vessels with pro-inflammatory macrophages (Anti(+) ≈ +0.45, #). The only significant difference between conditions was again Pro(+) versus Anti(−), and Anti(+) did not differ significantly from No Mac.(+). Taken together, anti-inflammatory macrophages limited both the loss of VE-Cadherin and the upregulation of ICAM-1, whereas pro-inflammatory macrophages amplified both.

### Macrophage polarization regulates vascular barrier permeability

To determine whether the opposing effects on endothelial protein expression in the presence of pro- versus anti-inflammatory macrophages produced a functional difference in barrier function, we measured the diffusive permeability coefficient (Pd) of the vessels. A 70-kDa FITC-dextran solution was perfused into the lumen and imaged by live confocal microscopy, and Pd was calculated as described previously (*30*, *32*) (Fig. 3A). Inflammatory stimulation of macrophage-free devices increased the permeability of the endothelial vessels (Fig. 3, B and C). Devices with pro-inflammatory macrophages had the steepest accumulation of FITC-dextran (Fig. 3C) and the highest Pd (median ≈ 6.5 × 10^−6^ cm/s), which was significantly higher than both the unstimulated macrophage-free control and the anti-inflammatory conditions (Fig. 3D). Devices with anti-inflammatory macrophages accumulated FITC-dextran at a rate comparable to unstimulated macrophage-free vessels (Fig. 3C) and had the lowest Pd (median ≈ 2.8 × 10^−6^ cm/s), approximately half the value measured in devices with pro-inflammatory macrophages (Fig. 3D). Stimulation increased Pd in macrophage-free vessels (median ≈ 2.7 to 4.3 × 10^−6^ cm/s) but not in vessels with anti-inflammatory macrophages, which remained at the unstimulated macrophage-free level whether or not they were stimulated. Neither anti-inflammatory condition differed significantly from the unstimulated macrophage-free control (P > 0.05). Macrophage polarization therefore regulates barrier function, with pro-inflammatory macrophages increasing vascular permeability and anti-inflammatory macrophages holding it at the unstimulated baseline.

**Fig. 3.**
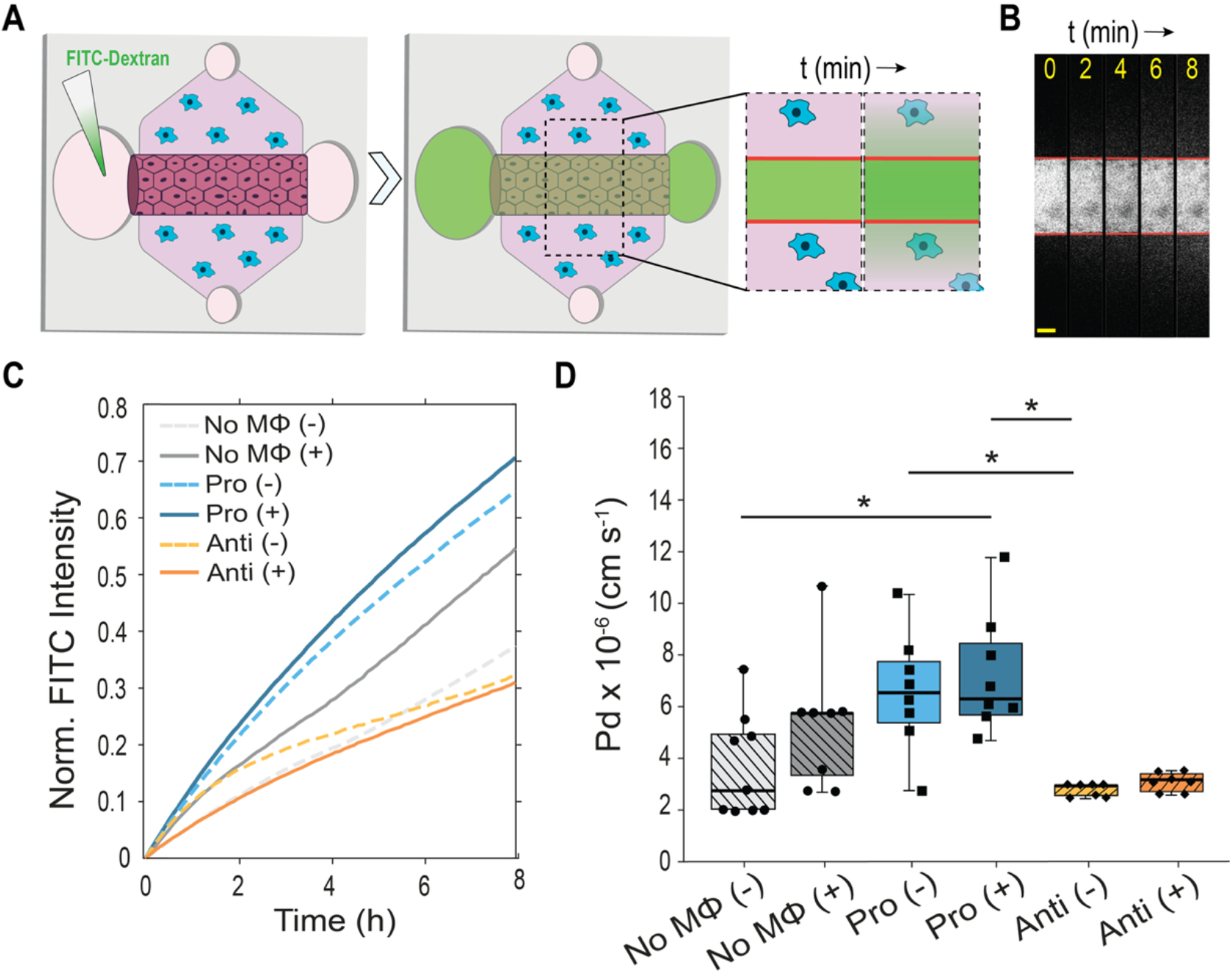
Macrophage polarization regulates vascular barrier permeability. (A) Schematic of the FITC-dextran permeability assay. A 70-kDa FITC-dextran solution is perfused into the HUVEC lumen and live-imaged at the vessel midplane by confocal microscopy, and the diffusive permeability coefficient Pd is computed as the slope of normalized extracellular matrix fluorescence accumulation over the 8-min window multiplied by the vessel-wall geometric factor (*30*). (B) Representative time-lapse confocal micrographs of FITC-dextran perfusion across the endothelial barrier at 2-min intervals (t = 0, 2, 4, 6 and 8 min). Red lines mark the vessel walls. Scale bar, 100 µm. (C) Condition mean of the normalized extracellular matrix fluorescence traces over the 8-min acquisition window. Solid lines, with stimulation. Dashed lines, without. (D) Diffusive permeability coefficient Pd (× 10^−6^ cm/s). All six conditions were acquired within each experimental series, across independent biological replicates. Kruskal–Wallis test with Bonferroni-corrected pairwise post hoc comparisons. *P < 0.05. N = 7–9, n = 2–3. In (D), the center line is the median, the box spans the interquartile range and the whiskers span the full range, with individual biological replicates overlaid. (+) denotes inflammatory stimulation and (−) its absence.

### Anti-inflammatory macrophages reduce neutrophil extravasation and migration distance

We next determined whether the differences in endothelial protein expression and barrier function between devices with pro- and anti-inflammatory macrophages differentially regulated the neutrophil response. We first quantified the magnitude of the response by counting the number of neutrophils that extravasated, a measure of how many cells gain access to the surrounding tissue. Neutrophil extravasation in response to a source of IL-8 and LTB_4_ was measured over 8 h (Fig. 4A). The pro-inflammatory and anti-inflammatory experiments were performed independently, each with its own macrophage-free control, and each macrophage condition was compared to its paired control. In the presence of anti-inflammatory macrophages, extravasation was significantly lower than macrophage-free devices at hours 5, 6, 7 and 8 (Fig. 4B). Extravasation in the presence of pro-inflammatory macrophages did not differ from its control at any timepoint (Fig. 4D). We then measured how far extravasated neutrophils migrated from the lumen edge over time, which reports how far into the tissue the responding cells travel, and compared each condition with its paired control. In the presence of anti-inflammatory macrophages, migration distance was significantly shorter than in macrophage-free devices at hours 2 through 7 (Fig. 4C). Migration distance in the presence of pro-inflammatory macrophages did not differ from its control at any timepoint (Fig. 4E). Anti-inflammatory macrophages therefore reduce both the number of neutrophils that cross the endothelium and the distance those neutrophils migrate into the extracellular matrix, whereas pro-inflammatory macrophages change neither.

**Fig. 4.**
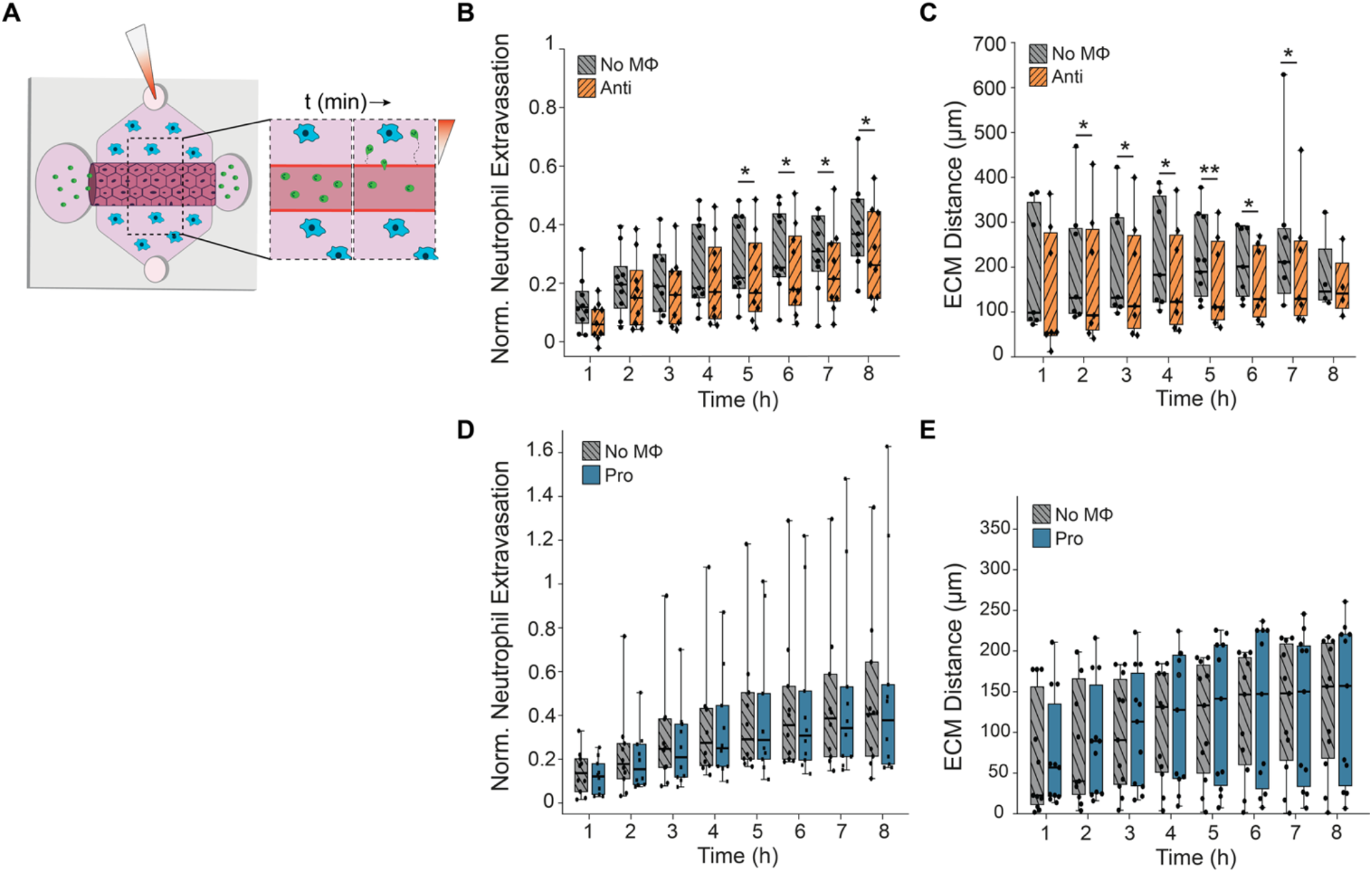
Anti-inflammatory macrophages reduce neutrophil extravasation and migration distance. (A) Schematic of the neutrophil extravasation assay. The TNF-α-primed HUVEC lumen is loaded with freshly isolated human neutrophils, a chemoattractant solution containing IL-8 and LTB_4_ is added to the top port, and confocal imaging tracks neutrophil extravasation and migration over 8 h. (B) Anti-inflammatory experiment, normalized neutrophil extravasation for anti-inflammatory devices versus their macrophage-free control. *P < 0.05. N = 9 paire(D) n = 2–3. (C) Anti-inflammatory experiment, neutrophil migration distance from the lumen edge. *P < 0.05, **P < 0.01. N = 8–9 paire(D) n = 2–3. (D) Pro-inflammatory experiment, normalized neutrophil extravasation for pro-inflammatory devices versus their macrophage-free control. P > 0.05. N = 10–11 paire(D) n = 2–3. (E) Pro-inflammatory experiment, neutrophil migration distance from the lumen edge. P > 0.05. N = 10–11 paire(D) n = 2–3. The pro-inflammatory and anti-inflammatory experiments were performed independently, each with its own macrophage-free control, and analyzed by within-experiment paired two-sample t-test at each timepoint without multiple-comparison correction. In (B) to (E), the center line is the median, the box spans the interquartile range and the whiskers span the full rang(E) with individual biological replicates overlaid. All devices received inflammatory stimulation.

One hypothesis for the reduced extravasation and shorter migration distance is the presence of anti-inflammatory macrophages reduces neutrophil motility. To test this, single-cell tracking was performed across the 8-h experimental window, and each macrophage condition was compared to its paired control. Speed, path length, straightness and directionality all declined over the 8-h window in both the anti-inflammatory devices and their macrophage-free controls (figs. S2 and S4), and the same metrics in the pro-inflammatory experiment are reported in figs. S2 and S3. No metric differed between devices with pro-inflammatory macrophages and their control at any timepoint. In the presence of anti-inflammatory macrophages, path length and speed were lower at hour 8, straightness at hour 5 and directionality at hour 2, with no differences at any other timepoint. Because the decline over the window occurs to the same extent in the macrophage-free controls, it does not account for the differences measured between conditions.

### Gas6 is most abundant in the presence of unstimulated anti-inflammatory macrophages

To identify factors that might contribute to the differences between devices with pro- and anti-inflammatory macrophages, we measured proteins secreted into the device microenvironment. Anti-inflammatory macrophages reduced neutrophil extravasation, whereas pro-inflammatory and macrophage-free devices did not differ from each other. We therefore looked for factors that had differing concentrations in devices with anti-inflammatory macrophages compared to the other two conditions. Conditioned media were collected from devices containing endothelial lumens with pro-inflammatory, anti-inflammatory, or no macrophages, before stimulation and 8 h after inflammatory stimulation. Five analytes were quantifiable across the panel (IL-6, CCL2, Gas6, CXCL1, and IL-1RA), and their concentrations were compared using two metrics, relative level and log_2_ fold change. Relative level is the position of each analyte within the six conditions, scaled from the lowest to the highest condition for that analyte, and shows where a factor is most and least abundant regardless of its absolute concentration (Fig. 5A). Log_2_ FC is the change in secreted concentration upon inflammatory stimulation, calculated for each replicate as its log_2_ concentration after stimulation minus the mean log_2_ concentration of the same condition without stimulation, and shows whether a factor increases or decreases with stimulation in a given macrophage environment (Fig. 5B). The absolute concentrations for each condition, unstimulated and 8 h after stimulation, are reported in fig. S5. These concentrations represent the amount present at a given timepoint and therefore reflect both secretion and consumption of each factor.

**Fig. 5.**
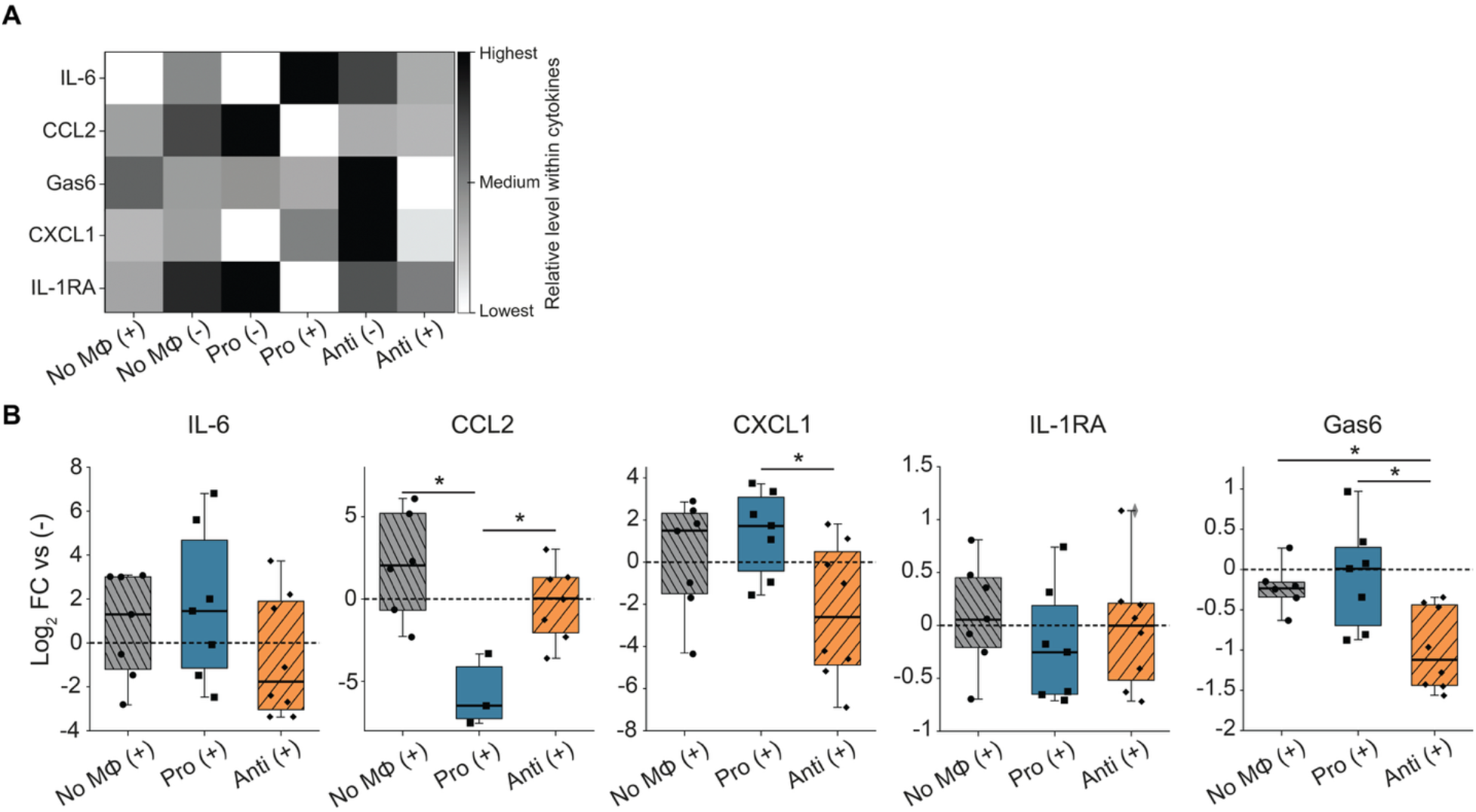
Macrophage phenotype shapes the cytokine profile of the inflammation-on-a-chip device. Conditioned media were collected from device lumens containing HUVECs with pro-inflammatory macrophages, antiinflammatory macrophages or no macrophages, at baseline and 8 h after inflammatory stimulation. (A) Heatmap of secreted IL-6, CCL2, Gas6, CXCL1 and IL-1RA across the six conditions, shaded from lowest to highest within each analyte. Shading is not comparable between analytes. (B) Stimulation-driven change for each of the five analytes in macrophage-free (No Mac), pro-inflammatory (Pro) and anti-inflammatory (Anti) conditions. For every analyte and condition, this change was calculated for each replicate as its log_2_ concentration after stimulation minus the mean log_2_ concentration of the same condition without stimulation (log_2_ FC). The dashed line marks no change (log_2_ FC = 0). One-way ANOVA with Tukey–Kramer post hoc correction. *P < 0.05. N = 3–8, n = 1, from a single MagPix measurement. In (B), the center line is the median, the box spans the interquartile range and the whiskers span the full range, with individual biological replicates overlaid. (+) denotes inflammatory stimulation and (−) its absence.

IL-6 was highest in devices with stimulated pro-inflammatory macrophages and next highest in devices with anti-inflammatory macrophages (Fig. 5A). However, the change in IL-6 with stimulation was greatest in devices with pro-inflammatory macrophages and lowest, falling below baseline, in devices with anti-inflammatory macrophages (Fig. 5B), although no comparison reached significance. CCL2 secretion decreased with stimulation (log_2_ FC < 0) in devices with pro-inflammatory macrophages and was significantly lower than in macrophage-free devices or devices with anti-inflammatory macrophages, which did not differ from each other (Fig. 5, A and B). CXCL1, a neutrophil chemoattractant, increased with stimulation (log_2_ FC > 0) in devices with pro-inflammatory macrophages and decreased (log_2_ FC < 0) in devices with anti-inflammatory macrophages, and the two conditions differed significantly (Fig. 5B). IL-1RA did not change with stimulation in any condition.

Gas6, the canonical AXL ligand, showed the clearest phenotype-linked pattern in the panel. Gas6 was highest in unstimulated devices with anti-inflammatory macrophages and lowest in those same devices after inflammatory stimulation (Fig. 5A). Gas6 also decreased with stimulation in devices with anti-inflammatory macrophages, significantly more than in macrophage-free or pro-inflammatory devices (Fig. 5B). Gas6 is therefore most abundant in the presence of anti-inflammatory macrophages before stimulation, and inflammatory stimulation reduces the concentration of Gas6 in the device. Of the analytes measured, Gas6 was the one that matched the pattern we were searching for.

### Gas6/AXL signaling mediates the reduced neutrophil response in the presence of anti-inflammatory macrophages

We found unstimulated devices with anti-inflammatory macrophages, had the highest Gas6 expression (Fig. 5A), the highest VE-Cadherin expression (Fig. 2B), the lowest ICAM-1 expression (Fig. 2C), the lowest Pd (Fig. 3, C and D), and the lowest neutrophil extravasation and migration (Fig. 4, B and C). We, therefore, directly tested whether Gas6/AXL signaling restricts the neutrophil response in the presence of anti-inflammatory macrophages. To inhibit signaling, devices were treated with the selective AXL receptor tyrosine kinase inhibitor bemcentinib (BGB324; 1 µM) throughout the neutrophil extravasation assay. We chose this concentration because it is well above the reported half-maximal inhibitory concentration and was still non-toxic to primary human leukocytes (*37*, *38*). Bemcentinib was included in both the lumen and the chemoattractant solution added to the top port, so that the inhibitor itself did not form a gradient across the matrix. Every biological replicate included four conditions: devices with anti-inflammatory macrophages and macrophage-free devices, each with and without bemcentinib.

AXL inhibition increased neutrophil extravasation both in devices with anti-inflammatory macrophages and in macrophage-free devices (Fig. 6A). In devices with anti-inflammatory macrophages, the increase was significant at every timepoint from hours 2 to 8. In macrophage-free devices, the increase became significant later, at hour 4, and continued through hour 8. At hour 4, mean normalized extravasation increased from 0.41 without inhibitor to 0.67 with inhibitor in macrophage-free devices, and from 0.31 to 0.43 in devices with anti-inflammatory macrophages. The effect was sustained, with no decay across the 8-h window in either condition. Migration distance was not significantly changed by AXL inhibition in either condition (Supplementary Fig. 6A). Without inhibitor, devices with anti-inflammatory macrophages had significantly shorter migration distances than macrophage-free controls at hours 4 and 6, and this difference was no longer significant with inhibitor, but the inhibitor-treated and untreated groups did not differ from one another at any timepoint. The reduced neutrophil extravasation measured in the presence of anti-inflammatory macrophages therefore depends, at least in part, on Gas6/AXL signaling, and can be reversed pharmacologically, even in the continued presence of anti-inflammatory macrophages. The increase in extravasation in macrophage-free devices indicates that AXL signaling also regulates the endothelium independently of macrophages.

**Fig. 6.**
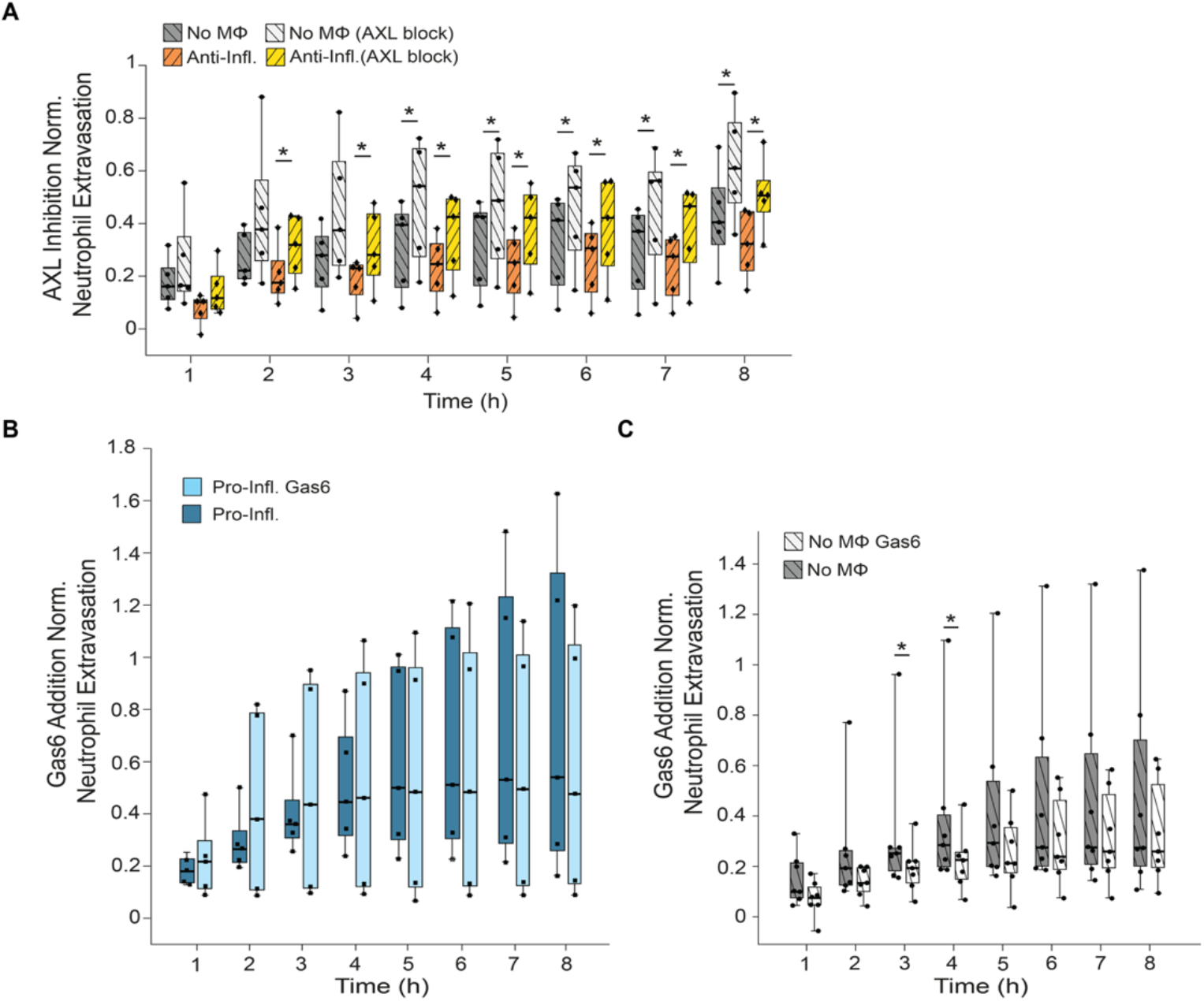
AXL inhibition increases neutrophil extravasation, whereas exogenous Gas6 reduces it in macrophage-free vessels but not in the presence of pro-inflammatory macrophages. (A) AXL inhibition, normalized neutrophil extravasation across four conditions, devices with anti-inflammatory macrophages (Anti-Infl.) and macrophage-free devices (No MΦ), each with or without the selective AXL inhibitor bemcentinib (1 µM, AXL block). Asterisks mark the drug comparison within each condition. Paired t-test at each timepoint. *P < 0.05. N = 5 paired, n = 2–3. (B) Gas6 addition to devices with pro-inflammatory macrophages (Pro-Infl.), with or without exogenous recombinant Gas6 (100 ng/mL), relative to the matched Gas6-free condition. Paired t-test at each timepoint. No timepoint reached significance (P > 0.05). N = 5 paired, n = 2–3. (C) Gas6 addition to macrophage-free devices (No MΦ), with or without exogenous recombinant Gas6 (100 ng/mL), relative to the matched Gas6-free condition. Because paired differences departed from normality at most timepoints (Shapiro–Wilk, P < 0.05, hours 2–8), conditions were compared by two-sided Wilcoxon signed-rank test at each timepoint. *P < 0.05 at hours 3 and 4. N = 7 paired (5 devices run alongside the pro-inflammatory-macrophage experiments in (B) plus 2 additional macrophage-free-only experiments), n = 2–5. In (A) to (C), the center line is the median, the box spans the interquartile range and the whiskers span the full range, with individual biological replicates overlaid. All devices received inflammatory stimulation.

We next performed the reverse experiment and determined whether exogenous Gas6 was sufficient to reduce neutrophil extravasation. Recombinant human Gas6 (100 ng/mL) (*39*, *40*) was added to the lumen and side ports of devices with pro-inflammatory macrophages and of macrophage-free devices, so that it reached both the endothelium and the surrounding matrix, and neutrophil extravasation was compared to matched Gas6-free controls. Gas6 significantly reduced extravasation in macrophage-free devices at hours 3 and 4 (Fig. 6C) but had no effect in devices with pro-inflammatory macrophages at any timepoint (Fig. 6B). At 100 ng/mL, the added Gas6 was far above the concentrations we measured in device-conditioned media (fig. S5), so the absence of a sustained effect does not reflect insufficient ligand. Migration distance did not differ between Gas6-supplemented and Gas6-free conditions at any timepoint (Supplementary Fig. 6B). Exogenous soluble Gas6 therefore restricts neutrophil extravasation in vessels without macrophages, but only at early timepoints, and does not reproduce the sustained reduction measured in the presence of anti-inflammatory macrophages.

## DISCUSSION

Our study identified anti-inflammatory macrophages as regulators of both endothelial activation and the neutrophil response to sterile inflammation using a three-dimensional inflammation-on-a-chip model developed using all primary human cells. Anti-inflammatory macrophages maintained endothelial barrier function and reduced neutrophil extravasation and migration into the surrounding matrix, whereas pro-inflammatory macrophages increased endothelial activation and permeability. Importantly, we identified Gas6/AXL signaling as a mediator of the reduced neutrophil extravasation measured in the presence of anti-inflammatory macrophages. Given the emerging role of Gas6 signaling in a range of diseases, including cancer, atherosclerosis, and sepsis, these results provide one potential mechanistic explanation for altered neutrophil recruitment in these settings.

Anti-inflammatory signals have been shown to alter endothelial barrier function, although the reported effects are not consistent. IL-10 reduces the adhesion of leukocytes to cytokine-activated human endothelial monolayers (41), and conditioned medium from IL-10-overexpressing macrophages restores transendothelial electrical resistance and limits dextran leakage across endothelial monolayers injured with lipopolysaccharide (42). In contrast, IL-4 increases permeability when applied directly to human endothelial cells (43). These studies measured barrier function in endothelial monolayers exposed to single cytokines. In microphysiological models, primary human synovial fibroblasts blunted a disease-like vascular phenotype when cultured with inflamed HUVECs (32), and, in vivo, perivascular macrophages protect vascular integrity (15). Our results with polarized primary human macrophages support the finding that anti-inflammatory macrophages improve vascular integrity. In our system, anti-inflammatory macrophages maintained VE-Cadherin expression, limited ICAM-1 upregulation and reduced diffusive permeability to approximately half the value measured in vessels with pro-inflammatory macrophages, indicating that the effect depends on polarization state rather than on the presence of macrophages. In contrast, the dominant effect of pro-inflammatory macrophages was an increase in ICAM-1 expression rather than a loss of VE-Cadherin, indicating that the endothelium was sensitized rather than physically disrupted. Increased ICAM-1 is expected to promote neutrophil capture by increasing avidity for β_2_-integrins (44, 45), and in vivo, ICAM-1 enrichment near tricellular endothelial junctions has been shown to mark preferred sites of leukocyte transmigration (46). Neutrophil extravasation in devices with pro-inflammatory macrophages did not exceed that of macrophage-free devices, which we interpret as our system being unable to resolve an increase above an already strong response rather than as evidence that endothelial sensitization has no effect. On the anti-inflammatory side, the reductions in extravasation and migration distance were larger and more sustained than the differences in single-cell motility, which were modest and confined to a few timepoints. Collectively, these results indicate that anti-inflammatory macrophages act primarily in regulating neutrophil extravasation, rather than regulating neutrophil migration once they are in the tissue. The gradual decline in motility seen across every condition is separate from this comparison, and follows from the chemoattractant being added once at the start of the assay, so that the IL-8 and LTB_4_ gradients flatten as both diffuse through the matrix.

Gas6 is a vitamin K-dependent ligand released by macrophages and other stromal cells that activates the TAM (TYRO3/AXL/MerTK) receptor tyrosine kinases on endothelial cells, where it has been linked to vascular homeostasis, junctional stability, and the limiting of leukocyte transmigration (39, 47–52). In our study, blocking AXL with bemcentinib (37, 38) increased neutrophil extravasation in devices with anti-inflammatory macrophages, reaching the same level of extravasation as macrophage-free conditions. Extravasation also increased in macrophage-free devices under AXL inhibition, indicating that endothelial Gas6/AXL signaling maintains a baseline barrier-protective tone, consistent with reports of autocrine AXL activity in resting endothelium (39, 53, 54). The Gas6/AXL signaling axis is better known elsewhere in immunity, where it sustains efferocytic, anti-inflammatory macrophage programs and supports the resolution of inflammation (55, 56). The axis also signals in chronic disease. In atherosclerosis it limits lesion progression by supporting the clearance of apoptotic cells within the plaque (57, 58), and in tumors it promotes an immunosuppressive microenvironment and is associated with resistance to therapy (59, 60). Our results indicate a role for Gas6 in acute, polarization-driven inflammation at the endothelium. Soluble Gas6 reduced extravasation when added to macrophage-free vessels, but only early in the response, and had no effect when pro-inflammatory macrophages were present. The manner in which Gas6 engages its receptor offers a possible explanation. Gas6’s Gla domain binds phosphatidylserine on cell membranes while its LG domains bind AXL, so Gas6 works by holding the receptor against a nearby membrane (47, 48). Therefore, it signals most effectively when anchored rather than free in solution.

Macrophages embedded in the matrix produce Gas6 continuously and release it alongside the membranes around them, whereas Gas6 added to the media is the same protein in an unanchored form. How the ligand is presented may, therefore, have just as strong an effect as its concentration. This also offers a way to reconcile diverging reports. Gas6 inhibits granulocyte adhesion to endothelial monolayers (53) and protects tight junctions during sepsis (40), yet has also been reported to promote leukocyte recruitment by enhancing interactions between endothelial cells, platelets and leukocytes (61), a route our platform excludes because it contains neither platelets nor flow. The Gas6/AXL axis is already being targeted clinically, with bemcentinib and related compounds in early-phase oncology trials (*37*), with stromal Gas6 blockade in pancreatic cancer shown to activate natural killer cells and reduce metastasis (*62*). That AXL blockade increased extravasation even in vessels without macrophages suggests such treatment could raise vascular permeability and neutrophil recruitment in tissues where excessive recruitment is itself harmful (*14*, *40*, *56*, *63*).

Several features of our model set limits on these conclusions. The platform is static and does not reproduce the wall shear stress that contributes to neutrophil capture in vivo. The three cell types are not donor-matched. The neutrophils and the monocytes come from separate healthy donors, and the HUVECs are a pooled commercial preparation, so the vessel is allogeneic to both leukocyte populations. Donor sex was neither recorded nor balanced under the approved protocol. Despite these constraints, the platform is well suited to investigate the questions these results raise. Because the macrophage compartment is built from primary human monocytes, it can be loaded with cells from patients to determine whether barrier-protective macrophage function is lost in conditions where neutrophil recruitment runs unchecked, such as systemic vasculitis or sepsis. More generally, because drugs can be delivered to defined compartments and their effects read out on both the vessel and the neutrophil response, the platform offers a route to testing candidate compounds in a fully human vascular environment.

Collectively, our results show that macrophage polarization plays an important regulatory role during sterile inflammation. Anti-inflammatory macrophages promote an environment in which the endothelium is less susceptible to inflammatory stimulation and fewer neutrophils cross into the surrounding tissue, whereas pro-inflammatory macrophages enhance endothelial activation. Gas6/AXL signaling is one mediator of this regulation. Because AXL inhibitors are already being given to patients, and because blocking AXL in our devices increased neutrophil extravasation even in vessels without macrophages, the macrophages surrounding a vessel deserve consideration both as determinants of vascular integrity and as a source of unintended vascular effects when this axis is targeted therapeutically.

## MATERIALS AND METHODS

### Ethics statement and human blood collection

All experimental procedures involving human blood were conducted in accordance with the Declaration of Helsinki and approved by the Institutional Review Board at the University of Colorado Boulder (Protocol #20-0082). Written informed consent was obtained from all healthy donors aged 18 to 65 years prior to blood collection. Whole blood was collected by venipuncture into K_2_EDTA tubes (BD Biosciences, 366643) and processed immediately.

### Sex and gender reporting

Healthy adult donors aged 18 to 65 years were enrolled without selection based on sex or gender. Donor sex was not recorded at the time of collection under the approved protocol, and the commercially sourced HUVEC preparation is pooled from multiple donors of unspecified sex. Sex- and gender-disaggregated analysis was therefore not possible, and sex was not considered as a biological variable in the analyses reported here. All findings should be interpreted as sex-unstratified; the potential influence of donor sex on macrophage polarization, neutrophil migration, endothelial barrier function and Gas6/AXL signaling is noted as a limitation in the Discussion.

### Isolation of primary human monocytes and neutrophils

Peripheral blood mononuclear cells were isolated by density-gradient centrifugation using Ficoll-Paque PLUS (Cytiva, 17144002). Whole blood was diluted 1:1 with sterile 1× PBS and layered over 15 mL of Ficoll-Paque PLUS in 50-mL conical tubes, then centrifuged at 400 × g for 35 min with soft deacceleration on. The mononuclear layer was collected, topped to 40 mL with PBS, and washed twice at 400 × g for 10 min, then twice at 100 × g for 10 min to remove platelets. Monocytes were isolated by negative selection using the Pan Monocyte Isolation Kit (Miltenyi Biotec, 130-096-537) and resuspended in pre-warmed AIM-V medium (Life Technologies, 12055091) at 1 × 10^6^ cells/mL. Peripheral blood neutrophils were isolated from fresh whole blood using the MACSxpress Neutrophil Isolation Kit (Miltenyi Biotec, 130-104-434), with residual erythrocytes depleted using the MACSxpress Erythrocyte Depletion Kit (Miltenyi Biotec, 130-098-196). Neutrophils were resuspended in warm EGM-2 and counted prior to use.

### Endothelial cell culture

Pooled primary human umbilical vein endothelial cells (HUVECs; PromoCell, C-12203 and 50305964) were maintained in Endothelial Growth Medium-2 (EGM-2; Lonza, NC9525043) in T-75 flasks between passages 2 and 7, with media refreshed every 2 days. At 80% confluence, cells were detached using 0.05% trypsin/0.02% EDTA (Fisher Scientific, 25300062) and reseeded at 3.75 × 10⁵ cells per flask.

### Inflammation-on-a-chip device fabrication and functionalization

Microfluidic devices were fabricated as previously described (*31*) and as employed in recent studies of neutrophil– endothelial transmigration (*34*, *35*) and synovial fibroblast–endothelial crosstalk(*32*). Top and bottom master molds were produced by stereolithography (Accura 5530 resin, Protolabs). Polydimethylsiloxane (PDMS; Sylgard 184, Dow) was cast in both molds and polymerized at 60 °C for 24 h. Cured PDMS layers were demolded and aligned, and a 0.337-mm-diameter PDMS rod was inserted between the layers to define the lumen geometry. The assembled device was bonded to a glass-bottom dish (MatTek, P50G-1.5-30-F) by oxygen plasma treatment. Devices were UV-sterilized for 15 min, then functionalized to promote hydrogel adhesion with 1% polyethylenimine (Sigma, 03880) for 10 min followed by 0.4% glutaraldehyde (Sigma, G6257) for 30 min, with three deionized-water washes.

### Device seeding and macrophage differentiation

Primary monocytes were embedded in neutralized type I rat-tail collagen (Corning, 354249) at 4 mg/mL supplemented with bovine plasma fibronectin (2 µg/mL; Fisher Scientific, NC0009247) and injected into the extracellular matrix ports. Devices were polymerized at room temperature for 15 min, transferred to 37 °C for 1.5 h, and PDMS rods were withdrawn to leave the lumen structure in place. Lumens were filled with AIM-V medium supplemented with M-CSF (2 ng/mL; PeproTech, 300-25). Day 0 denotes the timepoint 2 h after monocytes were embedded in the matrix. Monocytes were then differentiated in AIM-V with M-CSF for 5 days (Days 1 to 5) with daily media exchanges. On Day 6, macrophages were polarized for 24 h in medium that retained M-CSF (2 ng/mL): pro-inflammatory macrophages received IFN-γ (20 ng/mL; BioLegend) and LPS (100 ng/mL; Sigma, L4516); anti-inflammatory macrophages received IL-4 (20 ng/mL; Miltenyi Biotec, 130-093-915).

### HUVEC lumen seeding

On Day 7, lumens were washed three times with warm EGM-2 to remove polarization cytokines. HUVECs were resuspended in EGM-2 at 2 × 10^4^ cells/µL, introduced through the small side port, and devices were flipped every 15 min for 1 h to promote circumferential lumen attachment. Media was exchanged with fresh warm EGM-2 and devices were incubated overnight at 37 °C and 5% CO_2_, with EGM-2 exchanged twice daily thereafter.

### Cell viability assessment

Cell viability was assessed on Days 6, 7 and 9 using Calcein AM (1:200; Invitrogen, C3100MP), propidium iodide (1:1,000; Sigma, 53-706-05ML) and Hoechst 33342 (1 µg/mL; Thermo Fisher, H21486) in AIM-V. After 15 min at 37 °C, devices were washed three times with warm AIM-V, and three fields of view per device were captured on a Nikon A1R HD25 confocal microscope. Viability was quantified for pro-inflammatory and anti-inflammatory macrophages across three biological replicates. Each biological replicate was stained once, and the three fields acquired from that staining were treated as technical replicates and averaged to give one value per replicate. Viability is reported descriptively as the percentage of live cells per biological replicate (N = 3, n = 3); no significance test was applied, because viability remained uniformly high across conditions and timepoints and served as a quality-control check rather than a comparative endpoint.

### Macrophage phenotyping by immunofluorescence

Devices were fixed on Day 9 with 4% paraformaldehyde (Fisher Scientific, AC416780250) for 30 min. Surface staining was performed before permeabilization with FITC-anti-CD163 (1:500; Miltenyi Biotec, 130-099-969) and rabbit anti-CD80 (1:200; Life Technologies, PA5-85913) for 24 h at 4 °C. Cells were permeabilized with 0.3% Triton X-100 for 45 min and stained with APC-anti-CD68 (1:500; Miltenyi Biotec, 130-114-461) and Hoechst 33342 for 12 to 24 h at 4 °C; CD80 was detected with goat anti-rabbit IgG Alexa Fluor 594 (1:1,000; Thermo Fisher, A-11012). Confocal z-stacks were acquired with a 20× water-immersion objective. A custom MATLAB script segmented nuclei and calculated CD80^+^ and CD163^+^ percentages relative to total CD68^+^ cells. CD80^+^ and CD163^+^ percentages, and the Polarization Index (CD163%/CD80%), were compared between pro- and anti-inflammatory macrophages by Welch’s two-sample t-test on log_2_-transformed values.

### Validation of macrophage polarization by secreted markers

Validation of macrophage polarization by secreted markers. To confirm that the two polarization protocols produced functionally distinct populations, macrophages were profiled alone, without endothelial cells and without inflammatory stimulation, so that the measured analytes could be attributed to the macrophages themselves. Primary monocytes were embedded in collagen–fibronectin hydrogels in 96-well plates and differentiated and polarized as above. Hydrogels were washed three times with AIM-V and fresh AIM-V was added for a 24 h conditioning period. Conditioned media were collected and stored at −80 °C. A custom ProcartaPlex 5-plex Mix & Match immunoassay (Thermo Fisher) was run on a MagPix Luminex system (N = 4–5). The five analytes were chosen as polarization markers, pro-inflammatory IL-6 and TNF-α and anti-inflammatory IL-10, CD163 and MDC. For each replicate, an anti-inflammatory to pro-inflammatory ratio was computed as the sum of IL-10, CD163 and MDC divided by the sum of IL-6 and TNF-α. Individual analytes and the polarization ratio were log_2_-transformed and compared between pro- and anti-inflammatory macrophages by Welch’s two-sample t-test.

### Secretome profiling of the device microenvironment

To identify factors that could mediate the differences measured between conditions, the secretome was profiled in complete devices, with the endothelial lumen present and before and after inflammatory stimulation, so that the measurement reflects the microenvironment the neutrophils encounter rather than macrophage output alone. Conditioned media were collected from device lumens containing HUVECs with pro-inflammatory macrophages, anti-inflammatory macrophages or no macrophages (N = 3–8, n = 1) at baseline and 8 h after inflammatory stimulation. A custom ProcartaPlex Mix & Match immunoassay (Thermo Fisher) was run on a MagPix Luminex xMAP system. Five analytes were quantifiable and are reported here (IL-6, CXCL1, CCL2, Gas6 and IL-1RA); IL-1β and IL-10 were included in the panel but returned values at or below the lower limit of quantification in all wells and were therefore not analyzed. Concentrations were log_2_-transformed. Absolute concentrations were compared by Kruskal–Wallis with Dunn–Šidák post hoc correction, and stimulation-driven changes (log_2_ FC, each stimulated replicate relative to the mean of the same condition without stimulation) were compared by one-way ANOVA with Tukey–Kramer post hoc correction. Within each analyte and condition, log_2_ concentrations more than 1.5 interquartile ranges beyond the first or third quartile were excluded before analysis, applied only to groups with at least four values. For the heatmap, the mean log_2_ concentration of each condition was z-scored within each analyte and then scaled from lowest to highest across the six conditions, so shading reflects relative position within an analyte and is not comparable between analytes.

### Inflammatory stimulation

Unless stated otherwise, inflammatory stimulation means TNF-α (50 ng/mL; R&D Systems, 210-TA-020/CF) introduced into the lumen for 2 h at 37 °C followed by three EGM-2 washes, together with a chemoattractant solution of IL-8 (10 µM; Miltenyi Biotec, 130-122-353) and LTB_4_ (100 nM; Cayman Chemical, 20110) in the top port.

Macrophage viability, macrophage phenotyping and the secreted-factor profiling of macrophages in hydrogels were performed without inflammatory stimulation. The endothelial activation analysis used TNF-α alone, without the chemoattractant source.

### Endothelial activation analysis

For this experiment, stimulated conditions received TNF-α in the lumen for 2 h at 37 °C followed by three EGM-2 washes, without the chemoattractant source. On Day 9, endothelial lumens were fixed and stained with Hoechst 33342, Phalloidin-iFluor 594 (1:1,000; Abcam, ab176757), anti-VE-Cadherin (5:600; BD Biosciences, 561567) and anti-ICAM-1 (1:200; R&D Systems, BBA20). Confocal imaging used a 20× water-immersion objective. A custom MATLAB script generated a binary vessel mask by Gaussian smoothing and scaled Otsu thresholding; total fluorescence intensity within the mask was summed per image and normalized to nuclei count, then expressed as log_2_ FC relative to each replicate’s own unstimulated, macrophage-free condition. The five conditions (No Mac.(+), Pro(−), Pro(+), Anti(−) and Anti(+); throughout, (+) denotes this complete inflammatory stimulation, TNF-α with the IL-8 and LTB_4_ source, and (−) its absence) were analyzed by one-way ANOVA with Tukey–Kramer post hoc correction; each condition was additionally tested against baseline (log_2_ FC = 0) by one-sample t-test.

### Vascular barrier permeability assay

Vascular barrier permeability was assessed on Day 9 in six conditions (No Mac, Pro-inflammatory and Anti-inflammatory, each with or without inflammatory stimulation) using FITC-dextran perfusion followed by live confocal microscopy and quantification of the diffusive permeability coefficient (P_d_) as previously described^28,30^. TNF-α (50 ng/mL; R&D Systems, 210-TA-020/CF) was introduced into the lumen for 2 h at 37 °C, and a chemoattractant solution of IL-8 (10 µM; Miltenyi Biotec, 130-122-353) and LTB_4_ (100 nM; Cayman Chemical, 20110) was added to the top port, so that stimulated devices received the same complete stimulus used throughout. After 2 h, a 70-kDa FITC-dextran solution (125 µg/mL; Sigma, FD70S-100MG) in EGM-2 was introduced through the large port. Live confocal microscopy was performed at the vessel midplane with a 10× air objective, with images acquired every 6 s. A custom MATLAB script normalized the extracellular matrix signal to the initial lumen fluorescence, anchored it to zero at the first timepoint, fit a linear regression over the 8-min window, and computed P_d_ as the product of slope and a geometric factor derived from vessel geometry (vessel wall width / 4, expressed in cm), following the formulation established by Polacheck et al.(*30*). P_d_ values across all six conditions were compared by Kruskal–Wallis with Bonferroni-corrected pairwise post hoc comparisons.

### Neutrophil extravasation

Neutrophil extravasation was assessed in two independent experiments, a pro-inflammatory experiment comparing Pro devices to same-day macrophage-free controls and an anti-inflammatory experiment comparing Anti devices to same-day macrophage-free controls. The endothelial lumen was treated with TNF-α (50 ng/mL) for 2 h at 37 °C and washed three times with EGM-2. Freshly isolated neutrophils were resuspended in EGM-2 at 7.5 × 10^6^ cells/mL; half were labeled with Calcein AM (1:1,000) and recombined 1:1 with unstained neutrophils. The suspension was introduced into the lumen and 2 µL of IL-8 (10 µM) and LTB_4_ (100 nM) was loaded into the top port. Extravasation was monitored by confocal microscopy with a 10× air objective for 8 h, with acquisitions every 1 h. Extravasated neutrophils were counted within a 900 × 500-µm region of interest at the top of the lumen and normalized to the initial lumen loading density measured in a 400 × 500-µm central region at the start of imaging. Each device’s count was then baseline-subtracted against its own value at the first timepoint (timepoint 0), so that the metric reports the increase in extravasated neutrophils over each device’s starting count. We refer to this loading-normalized, timepoint-0–subtracted count as normalized extravasation from timepoint 0 (“normalized extravasation”) throughout. Within each device sheet, multipoints were assigned to the macrophage condition or to the same-day macrophage-free control, yielding one paired observation per biological replicate. Because the pro-inflammatory and anti-inflammatory experiments were performed on different days, each was analyzed independently and no cross-experiment comparison was performed. Within each experiment, the macrophage condition was compared to its matched macrophage-free control by paired two-sample t-test at every timepoint, without multiple-comparison correction.

### Neutrophil migration distance

Neutrophil migration distance was quantified from the same time-lapse acquisitions by measuring the distance of each neutrophil from the lumen edge, with raw pixel distances converted to µm using a calibration of 1.73 µm per pixel. As in the extravasation assay, each device provided both the macrophage condition and its matched macrophage-free control at separate imaging positions, and the positions belonging to each condition were averaged to a single biological-replicate value at each timepoint. The pro-inflammatory and anti-inflammatory experiments were analyzed independently using the same framework as the extravasation assay: paired two-sample t-test at every timepoint, without multiple-comparison correction.

### Neutrophil migration tracking

Single-cell tracking was performed with a 20× air objective in resonant scanning mode using the single-cell tracking module in NIS-Elements. Speed, path length, displacement and directional straightness (displacement / path length) were exported per device and processed with a custom MATLAB pipeline; directionality was additionally computed per biological replicate and timepoint bin. As for the extravasation and migration distance assays, the pro-inflammatory and anti-inflammatory experiments were analyzed independently, with each macrophage phenotype compared to its matched macrophage-free control by paired two-sample t-test at every timepoint.

### Pharmacological blockade of Gas6/AXL signaling

The selective AXL inhibitor bemcentinib (BGB324; 1 µM) was incorporated into the neutrophil extravasation assay. The 1 µM concentration was chosen because it lies roughly 70-fold above the reported half-maximal inhibitory concentration (~14 nM), ensuring near-complete AXL inhibition, while remaining below the ≥2.5 µM cytotoxicity threshold reported in normal primary human B, T and NK cells (*37*, *38*). Working inhibitor media (1 µM bemcentinib, 0.01% DMSO) was prepared fresh in EGM-2 daily. Devices were pre-incubated with working inhibitor media for 24 h before the assay, added to the lumen and to the top and bottom ports so that the inhibitor reached the extracellular matrix as well as the endothelium, and AXL was therefore blocked across the vessel and matrix before inflammatory stimulation. TNF-α was diluted in working inhibitor media and introduced into the lumen for 2 h at 37 °C; lumens were then washed three times with warm working inhibitor media. Neutrophils were resuspended in working inhibitor media, and the top-port chemoattractant solution was prepared in working inhibitor media to prevent a transmural drug gradient. Within each device, the twelve multipoints were assigned to four conditions: anti-inflammatory macrophages with bemcentinib (AXL-block), macrophage-free with bemcentinib (AXL-block), anti-inflammatory macrophages without bemcentinib, and macrophage-free without bemcentinib, with each device sheet contributing one paired observation per condition. Neutrophil loading, confocal acquisition and quantification of extravasation and migration distance followed the assays described above, with the inhibitor present in every solution. Two sets of comparisons were made by paired two-sample t-test at each timepoint (N = 5 paired biological replicates). The first tested the effect of the inhibitor within each vessel type, comparing macrophage-free devices with and without bemcentinib, and devices with anti-inflammatory macrophages with and without bemcentinib. The second tested the effect of the macrophages within each drug arm, comparing macrophage-free devices to devices with anti-inflammatory macrophages, separately in the untreated and the bemcentinib-treated groups.

### Exogenous Gas6 addition

To test whether Gas6 signaling is sufficient to restrict neutrophil extravasation, recombinant human Gas6 (R&D Systems, 885-GSB) was used in EGM-2 at a working concentration of 100 ng/mL, within the range previously shown to activate Gas6/AXL signaling in human endothelial cells (*39*, *40*). Gas6 was added to macrophage-free and pro-inflammatory macrophages devices, the two conditions in which Gas6 was least abundant (Fig. 5A), so that the experiment tested whether supplying the ligand to a low-Gas6 environment reproduced the reduced extravasation measured in the presence of anti-inflammatory macrophages. For 24 h, Gas6-supplemented EGM-2 (Gas6 (+)) or basal EGM-2 without Gas6 (Gas6 (−)) was added to the lumen and to the top and bottom ports, so that Gas6 reached the extracellular matrix as well as the endothelium. The lumen was then primed with TNF-α (50 ng/mL) prepared in the corresponding Gas6-supplemented or Gas6-free media for 2 h at 37 °C and washed three times with the same media. Freshly isolated neutrophils were resuspended in EGM-2 Gas6-supplemented or Gas6-free media, and a chemoattractant solution of IL-8 (10 µM) and LTB_4_ (100 nM) prepared in EGM-2 Gas6-supplemented or Gas6-free media was loaded into the top port, so that Gas6 was present throughout the assay in the Gas6 (+) condition, and neutrophil extravasation and migration distance from the lumen edge were imaged and quantified as described for the neutrophil extravasation and migration distance assays above. Within each device sheet, multipoints were assigned to four conditions (pro-inflammatory macrophages, pro-inflammatory macrophages with exogenous Gas6 (+), macrophage-free, and macrophage-free with exogenous Gas6 (+)), with each sheet contributing one paired observation per condition (N = 5 paired biological replicates). Two further experiments were run with macrophage-free devices only, giving N = 7 paired biological replicates for that condition. At each timepoint, the Gas6-supplemented condition was compared to its matched Gas6-free condition: by paired two-sample t-test for devices with pro-inflammatory macrophages, and, because the paired differences departed from normality (Shapiro–Wilk, P < 0.05, hours 2–8), by two-sided Wilcoxon signed-rank test for macrophage-free devices.

### Image acquisition

All microscopy was performed on a Nikon A1R HD25 laser scanning confocal microscope mounted on a Nikon Ti2-E inverted microscope system, controlled by NIS-Elements software. Cell viability, vascular barrier permeability, neutrophil extravasation and migration distance were imaged with a 10× air objective (NA 0.45). Single-cell neutrophil movement was imaged with a 20× air objective in resonant scanning mode. Macrophage phenotyping and HUVEC immunofluorescence were imaged with a 20× water-immersion objective in galvano scanning mode with sequential laser excitation. Image analysis was performed in NIS-Elements, Fiji (ImageJ) and custom MATLAB scripts.

### Statistics and reproducibility

All statistical analyses were performed in MATLAB R2024b (MathWorks). Each biological replicate (N) is an independent experiment performed with neutrophils and monocytes from a different donor, and replicate numbers therefore reflect the number of independent donor experiments completed for each condition rather than a predetermined sample size. Conditions that were run together within an experiment share that experiment, so N is matched within a comparison but can differ between comparisons; devices in which the endothelial lumen was incompletely formed, and analytes falling below the lower limit of quantification, were excluded, which reduces N for some conditions. Normality was assessed for every group by the Shapiro–Wilk test. Ratios, fold changes and analyte concentrations were right-skewed on the raw scale and were log_2_-transformed before parametric testing; for paired comparisons, normality was assessed on the within-replicate differences. Diffusive permeability and device-level cytokine concentrations remained non-normal after transformation and were analyzed non-parametrically. The statistical test applied to each readout, the number of biological (N) and technical (n) replicates and any post hoc correction are stated in the corresponding figure legend.

Box plots throughout show the median as the center line, the interquartile range as the box and the full range as the whiskers, with individual biological replicates overlaid. All P values derive from the tests named in the figure legends. A P value less than 0.05 was considered statistically significant.

## SUPPLEMENTARY MATERIALS

Figs. S1 to S6.

## ACKNOWLEDGMENTS

General

We thank the healthy blood donors who made this study possible, the staff of the BioFrontiers Advanced Light Microscopy Core at the University of Colorado Boulder for assistance with confocal imaging, and R. Kedl (University of Colorado Anschutz Medical Campus) for MagPix multiplex immunoassay support. We also thank members of the Hind Lab for discussion.

## Author contributions

A.I.-R. and L.E.H. conceived and designed the study. A.I.-R. performed the experiments, developed the image-analysis and statistical pipelines, and analyzed the data. R.S. contributed to the cell viability and vascular permeability experiments. A.I.-R. and L.E.H. interpreted the results and wrote the manuscript. L.E.H. supervised the project and acquired funding. All authors read and approved the final manuscript.

## Data and materials availability

Source data are provided with this paper. The MATLAB code and raw data files used to run the statistical tests and to generate the figures are deposited at Zenodo (https://doi.org/10.5281/zenodo.23166506)(64). Every group was checked for normality with a Shapiro–Wilk test before analysis, and this result was used to decide whether the data were analyzed with parametric or non-parametric tests. Raw confocal image datasets are available from the corresponding author on request.

---

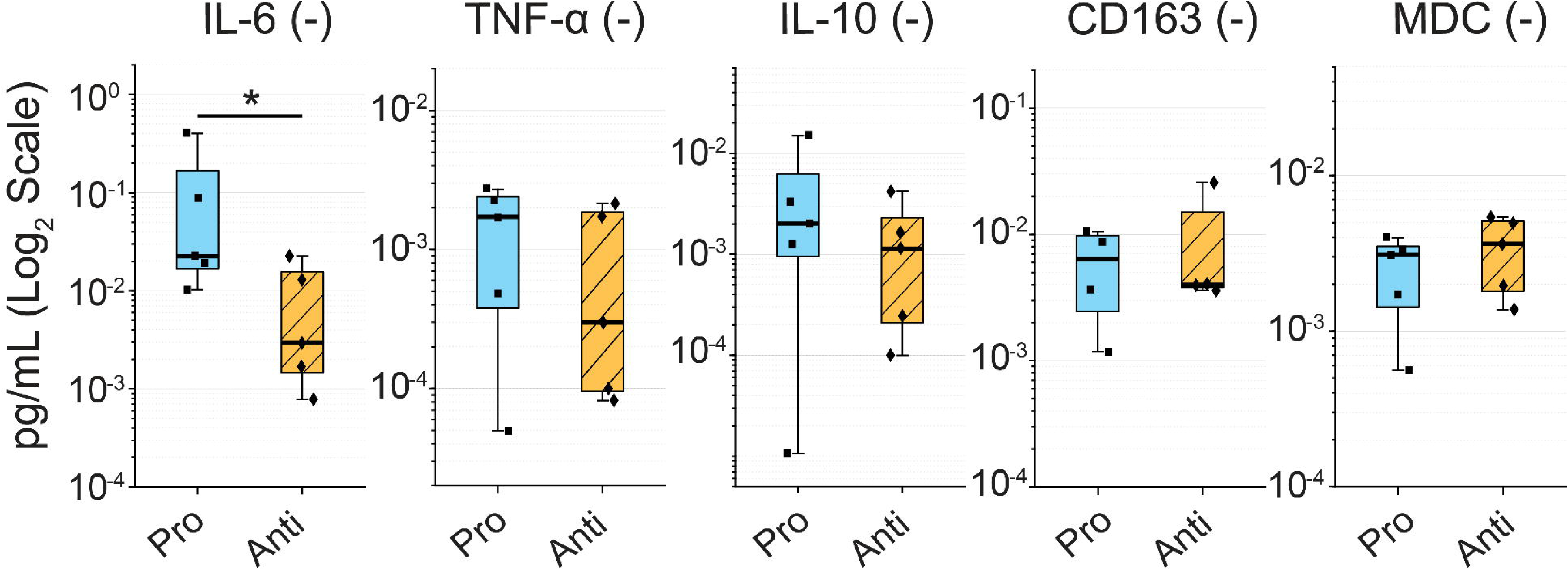

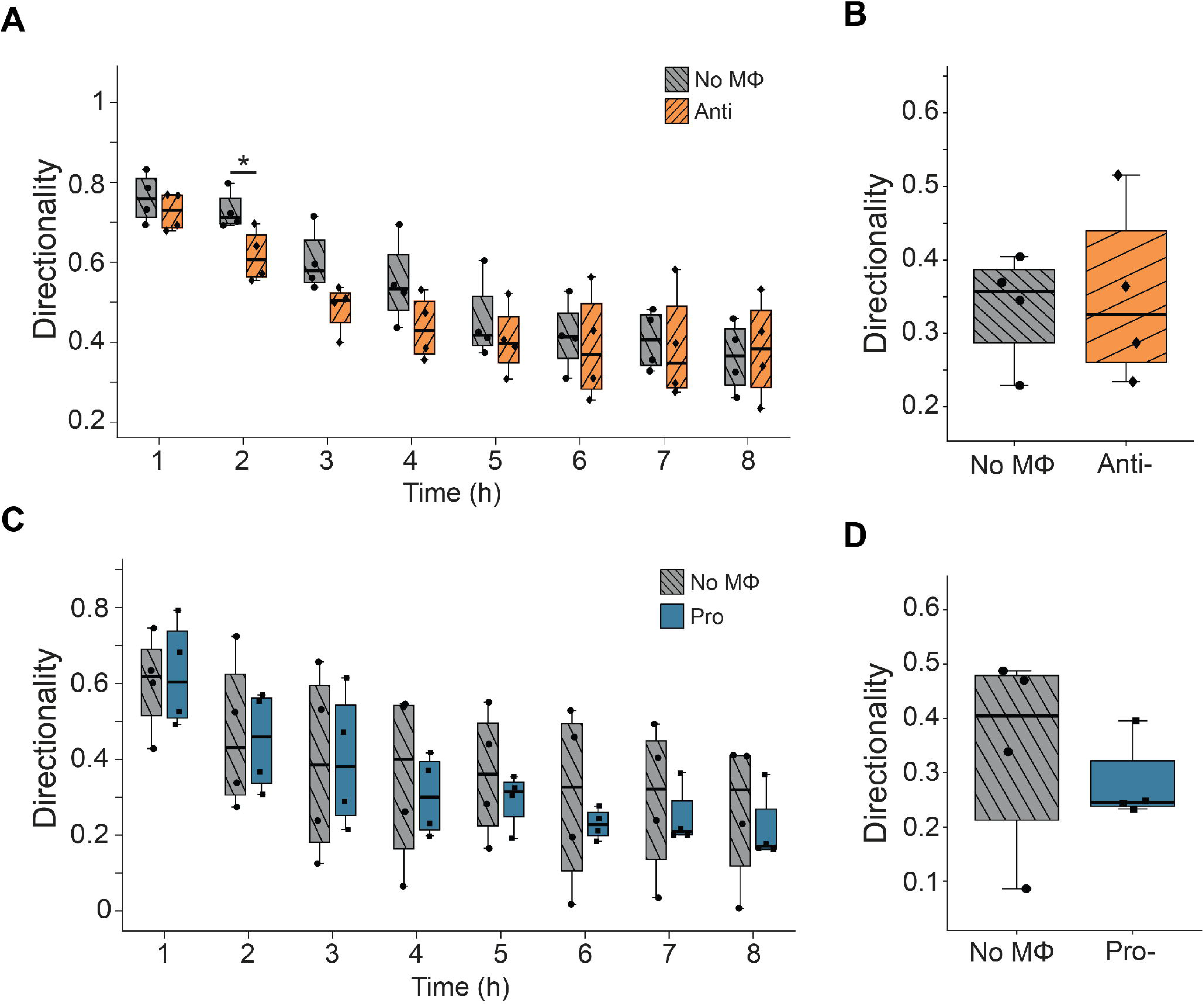

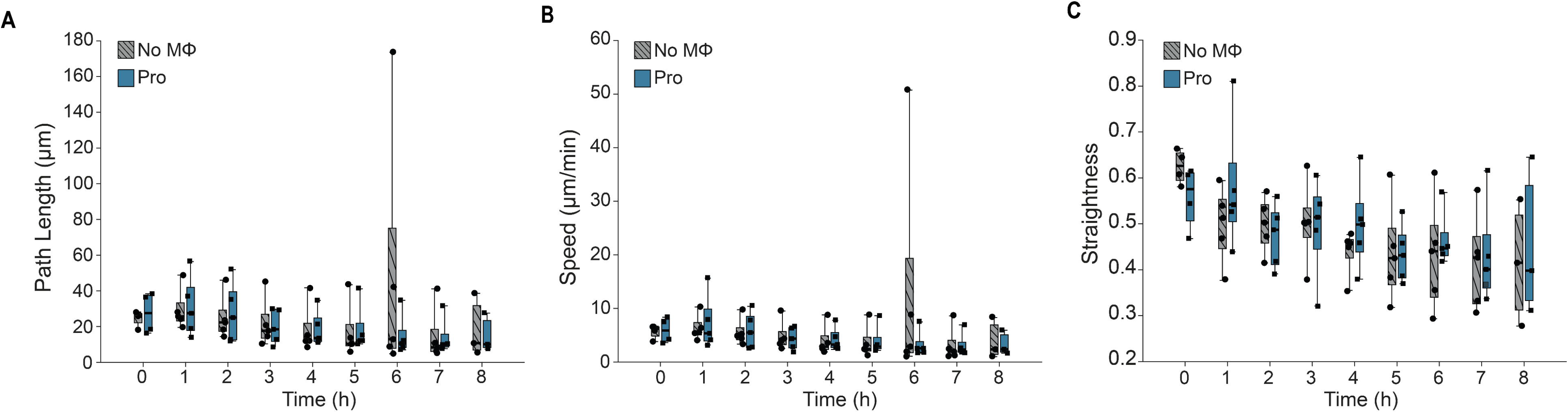

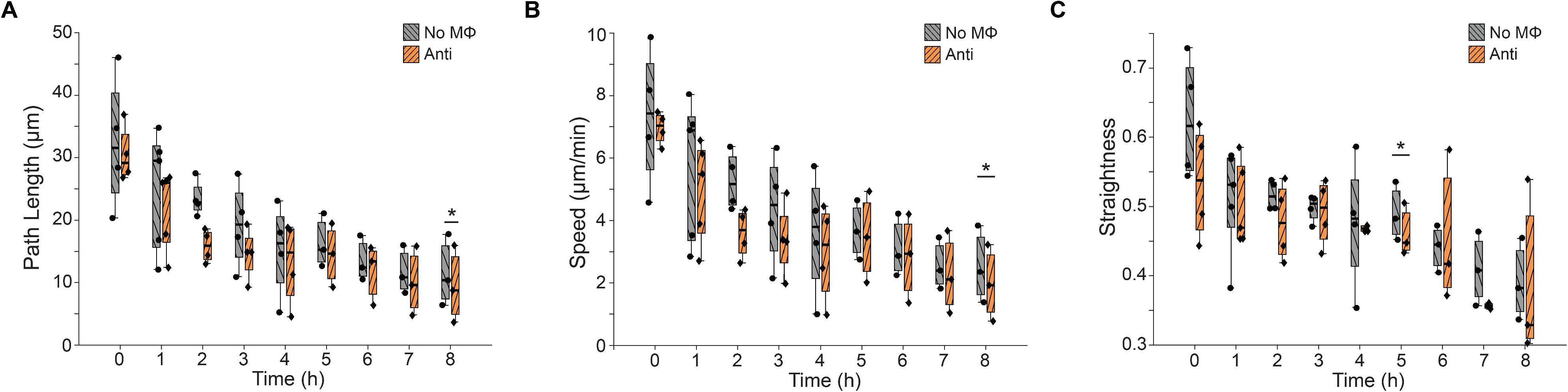

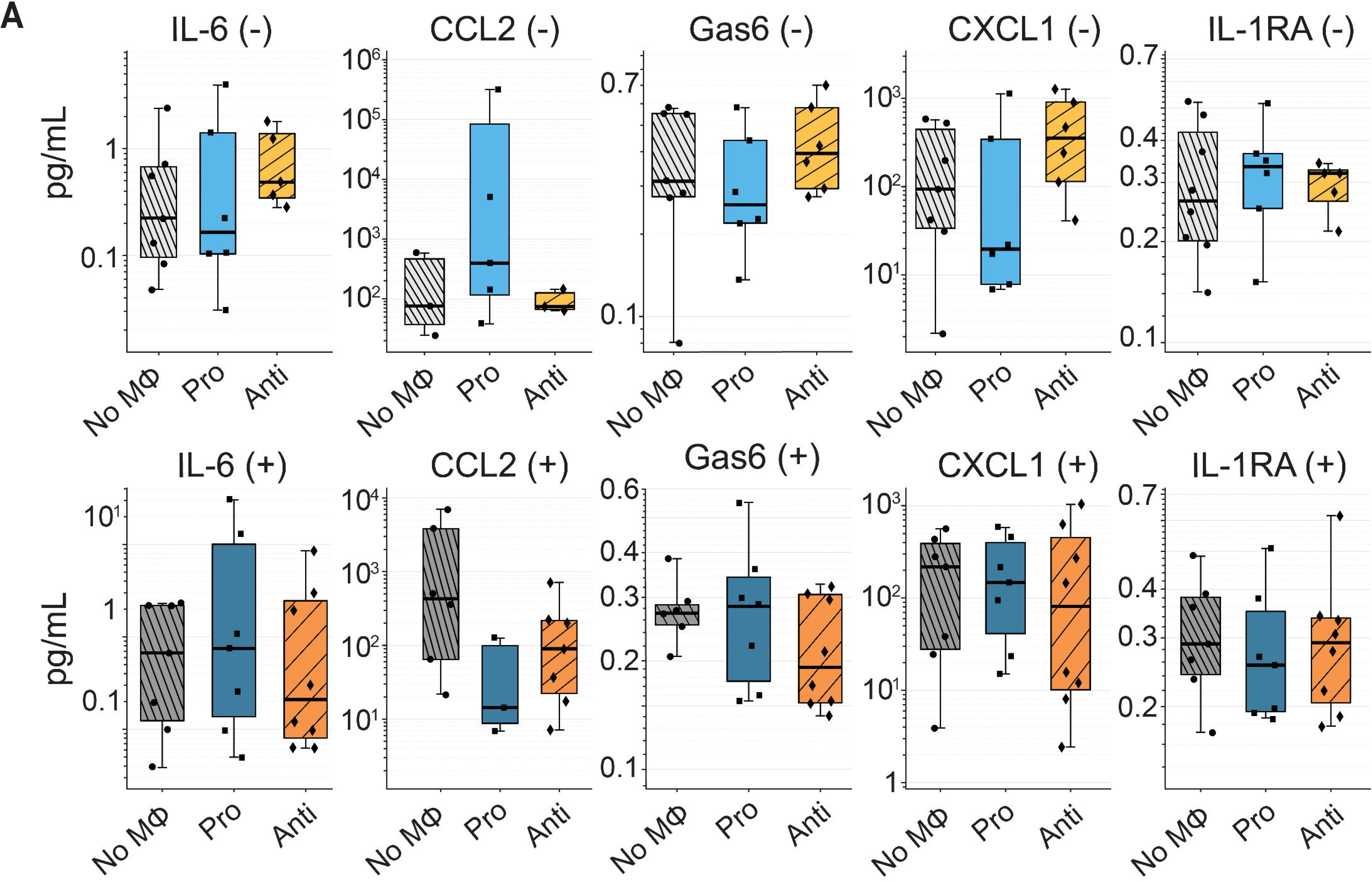

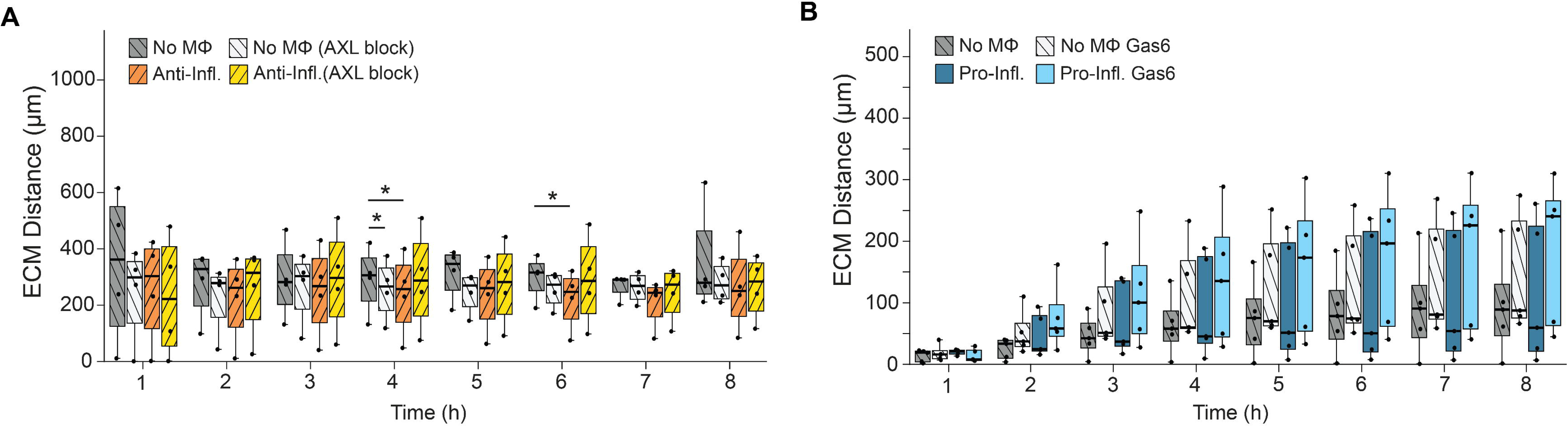

## REFERENCES AND NOTES

1. K. Ley, C. Laudanna, M. I. Cybulsky, S. Nourshargh, Getting to the site of inflammation: the leukocyte adhesion cascade updated. Nat. Rev. Immunol. 7, 678–689 (2007).

2. M. Phillipson, P. Kubes, The neutrophil in vascular inflammation. Nat. Med. 17, 1381– 1390 (2011).

3. W. A. Muller, Transendothelial migration: unifying principles from the endothelial perspective. Immunol. Rev. 273, 61–75 (2016).

4. E. Dejana, F. Orsenigo, M. G. Lampugnani, The role of VE-cadherin in vascular morphogenesis and permeability control. Prog. Mol. Biol. Transl. Sci. 116, 119–144 (2013).

5. D. Vestweber, How leukocytes cross the vascular endothelium. Nat. Rev. Immunol. 15, 692–704 (2015).

6. W. A. Muller, Getting leukocytes to the site of inflammation. Vet. Pathol. 50, 7–22 (2013).

7. K. Prame Kumar, A. J. Nicholls, C. H. Y. Wong, Partners in crime: neutrophils and monocytes/macrophages in inflammation and disease. Cell Tissue Res. 371, 551–565 (2018).

8. E. Anitua, M. Troya, M. H. Alkhraisat, Dialogues in immunity: the interplay between neutrophils and macrophages. Biomedicines 14, 520 (2026).

9. A. Lapenna, M. De Palma, C. E. Lewis, Perivascular macrophages in health and disease. Nat. Rev. Immunol. 18, 689–702 (2018).

10. A. Abtin, Perivascular macrophages mediate neutrophil recruitment during bacterial skin infection. Nat. Immunol. 15, 45–53 (2014).

11. H. He, Perivascular macrophages limit permeability. Arterioscler. Thromb. Vasc. Biol. 36, 2203–2212 (2016).

12. X. Ren, Macrophage–endothelial cell crosstalk orchestrates neutrophil recruitment in inflamed mucosa. J. Clin. Invest. 133, e170733 (2023).

13. W. Li, Heart-resident CCR2+ macrophages promote neutrophil extravasation through TLR9/MyD88/CXCL5 signaling. JCI Insight 1, e87315 (2016).

14. S. Osorio-Valencia, B. Zhou, Roles of macrophages and endothelial cells and their crosstalk in acute lung injury. Biomedicines 12, 632 (2024).

15. B. Weijts, J. A. A. Demmers, C. Robin, Blood vessel-resident macrophages safeguard blood and vessel integrity in zebrafish. Nat. Immunol., doi: 10.1038/s41590-026-02481-y (2026).

16. P. J. Murray, Macrophage activation and polarization: nomenclature and experimental guidelines. Immunity 41, 14–20 (2014).

17. A. Sica, A. Mantovani, Macrophage plasticity and polarization: in vivo veritas. J. Clin. Invest. 122, 787–795 (2012).

18. A. Shapouri-Moghaddam, Macrophage plasticity, polarization, and function in health and disease. J. Cell. Physiol. 233, 6425–6440 (2018).

19. S. K. Moestrup, H. J. Møller, CD163: a regulated hemoglobin scavenger receptor with a role in the anti-inflammatory response. Ann. Med. 36, 347–354 (2004).

20. E. Hickman, Expanded characterization of in vitro polarized M0, M1, and M2 human monocyte-derived macrophages. PLoS ONE 18, e0279037 (2023).

21. B. McDonald, Intravascular danger signals guide neutrophils to sites of sterile inflammation. Science 330, 362–366 (2010).

22. J. Mestas, C. C. W. Hughes, Of Mice and Not Men: Differences between Mouse and Human Immunology. J. Immunol., doi: 10.4049/jimmunol.172.5.2731 (2004).

23. W. M. Nauseef, Human neutrophils ≠ murine neutrophils: Does it matter? Immunol. Rev. 314, 442–456 (2023).

24. N. D. Tan, D. Davidson, Comparative differences and combined effects of interleukin-8, leukotriene B4, and platelet-activating factor on neutrophil chemotaxis of the newborn. Pediatr. Res. 38, 11–16 (1995).

25. F. Lin, Neutrophil migration in opposing chemoattractant gradients using microfluidic chemotaxis devices. Ann. Biomed. Eng. 33, 475–482 (2005).

26. P. V. Afonso, LTB4 is a signal-relay molecule during neutrophil chemotaxis. Dev. Cell 22, 1079–1091 (2012).

27. T. Lämmermann, Neutrophil swarms require LTB4 and integrins at sites of cell death in vivo. Nature 498, 371–375 (2013).

28. D. E. Ingber, Human organs-on-chips for disease modelling, drug development and personalized medicine. Nat. Rev. Genet. 23, 467–491 (2022).

29. M. Humayun, Innate immune cell response to host–parasite interaction in a human intestinal tissue microphysiological system. Sci. Adv. 8, eabm8012 (2022).

30. W. J. Polacheck, M. L. Kutys, J. B. Tefft, C. S. Chen, Microfabricated blood vessels for modeling the vascular transport barrier. Nat. Protoc. 14, 1425–1454 (2019).

31. J. A. Jiménez-Torres, S. L. Peery, K. E. Sung, D. J. Beebe, LumeNEXT: a practical method to pattern luminal structures in ECM gels. Adv. Healthc. Mater. 5, 198–204 (2016).

32. H. M. Zlotnick, Synovial fibroblasts modulate endothelial activation in an acute injury on a chip model. Proc. Natl. Acad. Sci. U. S. A. 123, e2524677123 (2026).

33. H. Kim, Macrophages-triggered sequential remodeling of endothelium–interstitial matrix to form pre-metastatic niche in microfluidic tumor microenvironment. Adv. Sci. 6, 1900195 (2019).

34. C. J. Calo, T. Patil, M. Palizzi, N. Wheeler, L. E. Hind, Collagen concentration regulates neutrophil extravasation and migration in response to infection in an endothelium dependent manner. Front. Immunol. 15, 1405364 (2024).

35. A. Ignes-Romeu, H. K. Weppner, T. Kaur, M. Singh, L. E. Hind, THP-1 Macrophages Limit Neutrophil Transendothelial Migration in a Model Infection. Cell. Mol. Bioeng. 17, 279–293 (2024).

36. H. K. Weppner, M. A. Singh, E. N. Katsnelson, D. J. Douin, M. D. McCarter, R. P. Tobin, L. E. Hind, Human M-MDSCs impair neutrophil migration in the infectious microenvironment. J. Leukoc. Biol. 117, qiaf165 (2025).

37. S. J. Holland, R428, a selective small molecule inhibitor of Axl kinase, blocks tumor spread and prolongs survival in models of metastatic breast cancer. Cancer Res. 70, 1544–1554 (2010).

38. A. K. Ghosh, The novel receptor tyrosine kinase Axl is constitutively active in B-cell chronic lymphocytic leukemia and acts as a docking site of nonreceptor kinases. Blood 117, 1928–1937 (2011).

39. J. G. Ganopolsky, M. R. Abid, W. C. Aird, M. D. Blostein, GAS6-induced signaling in human endothelial cells is mediated by FOXO1a. J. Thromb. Haemost. 6, 1804–1811 (2008).

40. J. Ni, Gas6 attenuates sepsis-induced tight junction injury and vascular endothelial hyperpermeability via the Axl/NF-κB signaling pathway. Front. Pharmacol. 10, 662 (2019).

41. T. Krakauer, IL-10 inhibits the adhesion of leukocytic cells to IL-1-activated human endothelial cells. Immunol. Lett. 45, 61–65 (1995).

42. L. Yi, T. Weng, P. Nie, L. Zhu, M. Gao, H. Jia, S. Yang, X. Li, L. Zhang, Y. Xu, P. Ma, M. Hu, Overexpression of interleukin-10 in engineered macrophages protects endothelial cells against LPS-induced injury in vitro. FEBS Open Bio 12, 605–615 (2022).

43. T. Skaria, J. Burgener, E. Bachli, G. Schoedon, IL-4 causes hyperpermeability of vascular endothelial cells through Wnt5A signaling. PLoS ONE 11, e0156002 (2016).

44. M. J. Allingham, J. D. van Buul, K. Burridge, ICAM-1-mediated, Src- and Pyk2-dependent vascular endothelial cadherin tyrosine phosphorylation is required for leukocyte transendothelial migration. J. Immunol. 179, 4053–4064 (2007).

45. L. Yang, ICAM-1 regulates neutrophil adhesion and transcellular migration of TNF-α-activated vascular endothelium under flow. Blood 106, 584–592 (2005).

46. R. Sumagin, E. Lomakina, I. H. Sarelius, Intercellular adhesion molecule-1 enrichment near tricellular endothelial junctions is preferentially associated with leukocyte transmigration. J. Immunol. 184, 5242–5252 (2010).

47. G. Lemke, Biology of the TAM receptors. Cold Spring Harb. Perspect. Biol. 5, a009076 (2013).

48. C. V. Rothlin, E. A. Carrera-Silva, L. Bosurgi, S. Ghosh, TAM receptor signaling in immune homeostasis. Annu. Rev. Immunol. 33, 355–391 (2015).

49. F. Li, L. Xu, C. Li, F. Hu, Y. Su, Immunological role of Gas6/TAM signaling in hemostasis and thrombosis. Thromb. Res. 238, 161–171 (2024).

50. K. E. Happonen, S. Tran, M. Mörgelin, R. Prince, S. Calzavarini, A. Angelillo-Scherrer, B. Dahlbäck, The Gas6-Axl Protein Interaction Mediates Endothelial Uptake of Platelet Microparticles. J. Biol. Chem. 291, 10586–10601 (2016).

51. G. Lemke, C. V. Rothlin, Immunobiology of the TAM receptors. Nat. Rev. Immunol. 8, 327–336 (2008).

52. M. Gallicchio, Inhibition of vascular endothelial growth factor receptor 2-mediated endothelial cell activation by Axl tyrosine kinase receptor. Blood 105, 1970–1976 (2005).

53. G. C. Avanzi, GAS6 inhibits granulocyte adhesion to endothelial cells. Blood 91, 2334– 2340 (1998).

54. V. A. Korshunov, Axl-dependent signalling: a clinical update. Clin. Sci. 122, 361–368 (2012).

55. L. Jiménez-García, The TAM receptor tyrosine kinases Axl and Mer drive the maintenance of highly phagocytic macrophages. Front. Immunol. 13, 960401 (2022).

56. Y. Tang, H. Hu, Q. Xie, J. Shen, GAS6/AXL signaling promotes M2 microglia efferocytosis to alleviate neuroinflammation in sepsis-associated encephalopathy. Cell Death Discov. 11, 268 (2025).

57. E. Lutgens, Genetic loss of Gas6 induces plaque stability in experimental atherosclerosis. J. Pathol. 216, 55–63 (2008).

58. T. Prouse, S. Majumder, R. Majumder, Functions of TAM receptors and ligands Protein S and Gas6 in atherosclerosis and cardiovascular disease. Int. J. Mol. Sci. 25, 12736 (2024).

59. M. Tanaka, D. W. Siemann, Gas6/Axl signaling pathway in the tumor immune microenvironment. Cancers 12, 1850 (2020).

60. X. Zhai, Gas6/AXL pathway: immunological landscape and therapeutic potential. Front. Oncol. 13, 1121130 (2023).

61. M. Tjwa, Gas6 promotes inflammation by enhancing interactions between endothelial cells, platelets, and leukocytes. Blood 111, 4096–4105 (2008).

62. L. Ireland, T. Luckett, M. C. Schmid, A. Mielgo, Blockade of stromal Gas6 alters cancer cell plasticity, activates NK cells, and inhibits pancreatic cancer metastasis. Front. Immunol. 11, 297 (2020).

63. M. Rizzi, S. Tonello, D. D’Onghia, P. P. Sainaghi, Gas6/TAM axis involvement in modulating inflammation and fibrosis in COVID-19 patients. Int. J. Mol. Sci. 24, 951 (2023).

64. A. A. Ignes-Romeu, R. Sood, L. Hind, Data and code for: Anti-inflammatory macrophages restrict neutrophil extravasation through Gas6/AXL signaling and maintain endothelial barrier function; 10.5281/zenodo.23166506.

