## Supplemental Data for "Anti-inflammatory macrophages limit endothelial inflammation and restrict neutrophil recruitment through Gas6/AXL signaling"

### SUPPLEMENTARY FIGURE LEGENDS

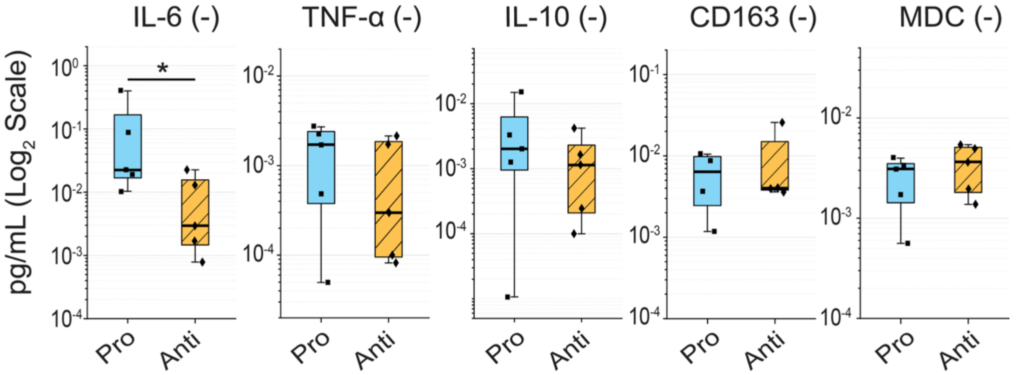


**Fig. S1. Absolute cytokine concentrations in the inflammation-on-a-chip ECM.** Absolute secreted concentrations (pg/mL, $\log_2$ scale) of the two pro-inflammatory analytes (IL-6 and TNF-α) and the three anti-inflammatory analytes (IL-10, soluble CD163 and MDC/CCL22), measured by multiplex MagPix Luminex assay in media conditioned by pro-inflammatory (Pro) and anti-inflammatory (Anti) macrophages embedded in the 3D collagen–fibronectin extracellular matrix. These are the raw values underlying the Anti/Pro secretion ratio in Fig. 1d. Welch's t-test on $\log_2$-transformed values. IL-6 was significantly higher in pro-inflammatory macrophages (*$P < 0.05$) and no other analyte differed significantly. N = 4–5, n = 1. The center line is the median, the box spans the interquartile range and the whiskers span the full range, with individual biological replicates overlaid. No inflammatory stimulation was applied.

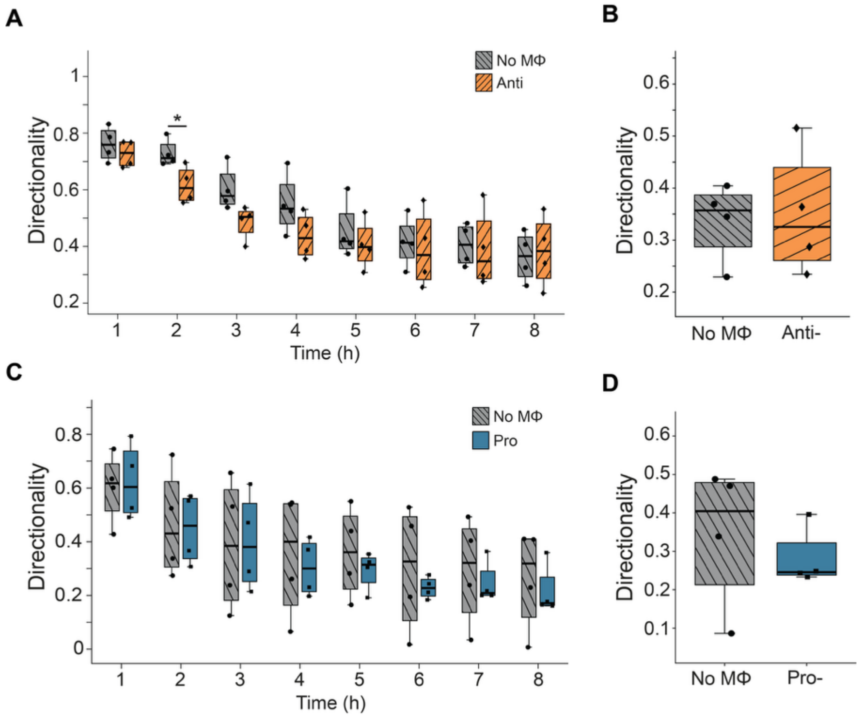


**Fig. S2. Neutrophil directionality over the 8-h window.** (A) Anti-inflammatory experiment, directionality (net displacement divided by total path length) for the anti-inflammatory device versus its matched macrophage-free control over time. *$P < 0.05$, with no significant difference at other timepoints. (B) Anti-inflammatory experiment, time-collapsed summary. Not significant. (C) Pro-inflammatory experiment, directionality for the pro-inflammatory device versus its matched control. P > 0.05. (D) Pro-inflammatory experiment, time-collapsed summary. Not significant. Paired two-sample t-test at each timepoint. N = 4, n = 2–3. The center line is the median, the box spans the interquartile range and the whiskers span the full range, with individual biological replicates overlaid. All devices received inflammatory stimulation.

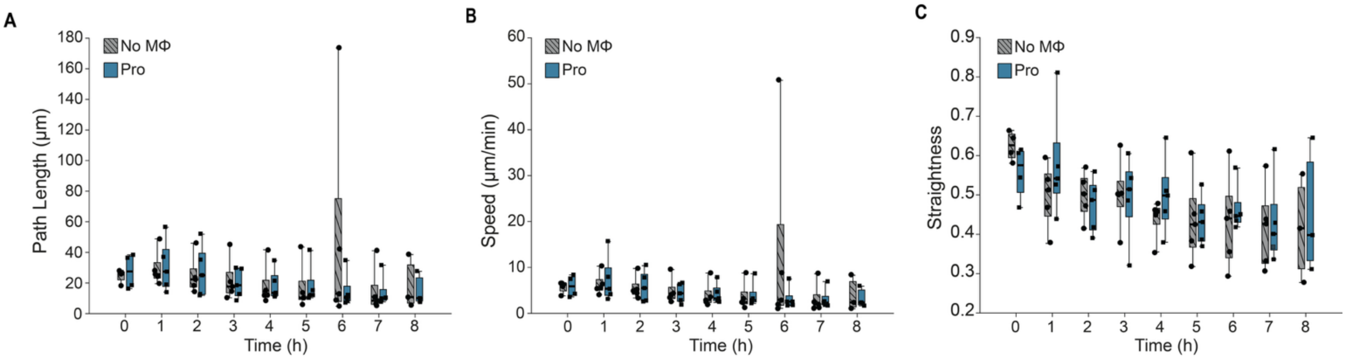


**Fig. S3. Neutrophil movement in the pro-inflammatory experiment over the 8-h window.** (A) Per-interval path length (µm). (B) Instantaneous speed (µm/min). (C) Directional straightness (displacement divided by path length). Pro-inflammatory device versus matched macrophage-free control, paired two-sample t-test at each timepoint. No timepoint reached significance. N = 4, n = 2–3. The center line is the median, the box spans the interquartile range and the whiskers span the full range, with individual biological replicates overlaid. All devices received inflammatory stimulation.

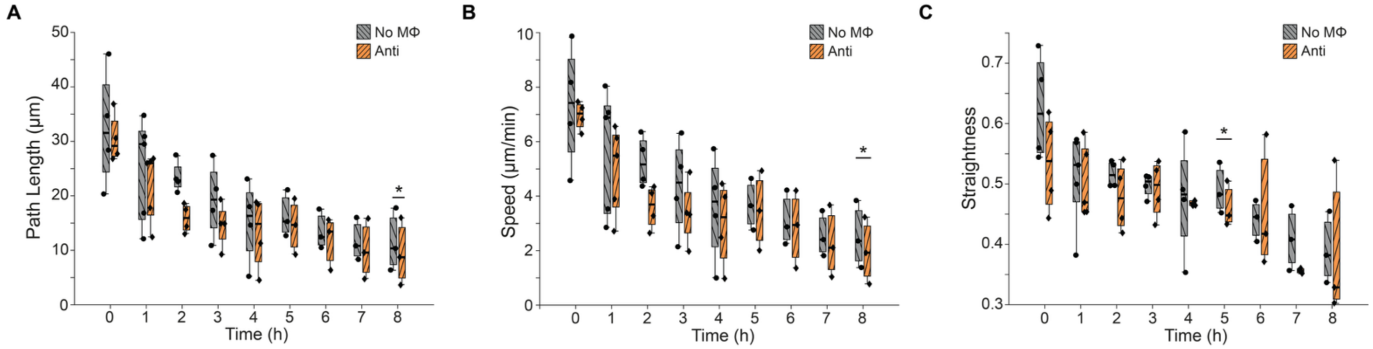


**Fig. S4. Neutrophil movement in the anti-inflammatory experiment over the 8-h window.** (A) Per-interval path length (µm). (B) Instantaneous speed (µm/min). (C) Directional straightness (displacement divided by path length). Anti-inflammatory device versus matched macrophage-free control, paired two-sample t-test at each timepoint. *$P < 0.05$. N = 4, n = 2–3. The center line is the median, the box spans the interquartile range and the whiskers span the full range, with individual biological replicates overlaid. All devices received inflammatory stimulation.

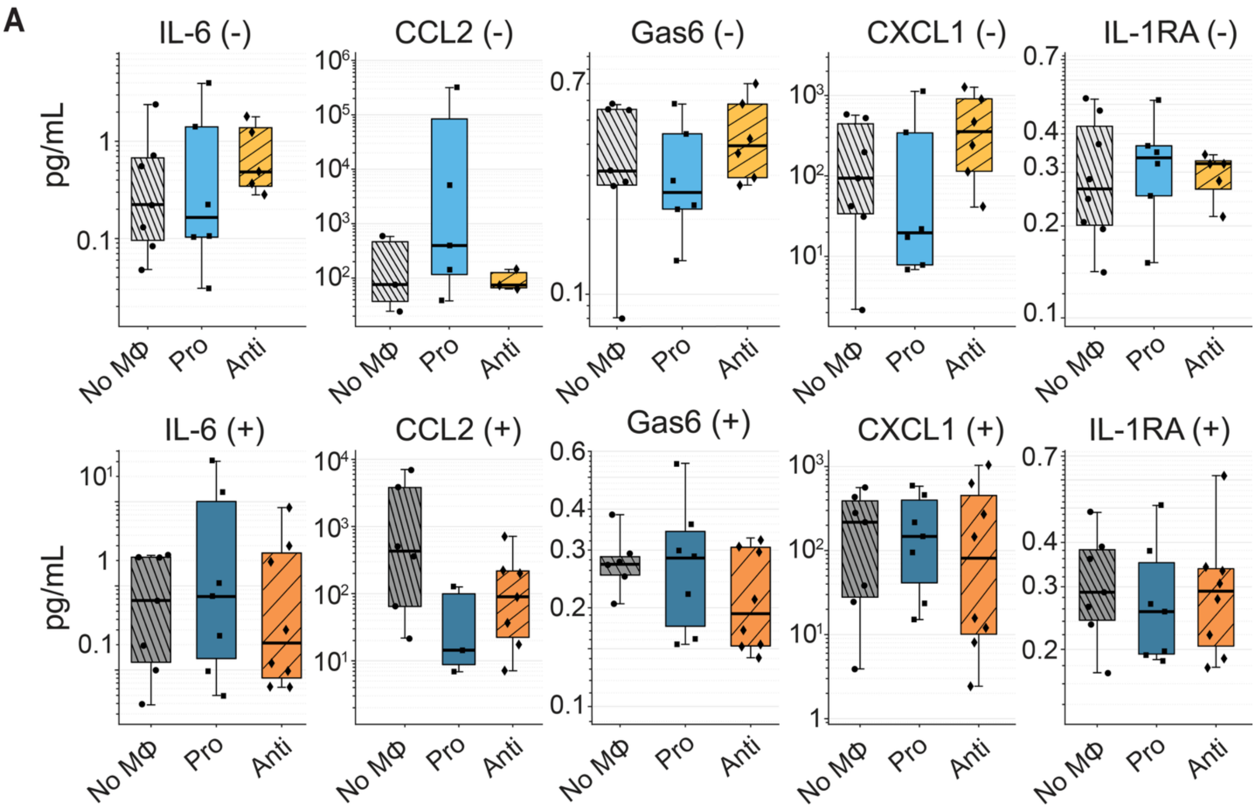


**Fig. S5. Device conditioned-media cytokine concentrations.** Absolute secreted concentrations (pg/mL, log scale) of the five quantifiable analytes (IL-6, CCL2, Gas6, CXCL1 and IL-1RA) measured in the device lumen for macrophage-free (No MΦ), pro-inflammatory (Pro) and anti-inflammatory (Anti) conditions, without inflammatory stimulation (top row) and 8 h after stimulation (bottom row). Concentrations were $log_2$-transformed and compared by Kruskal–Wallis with Dunn–Šidák post hoc correction. N = 3–8, n = 1, from a single MagPix measurement. The center line is the median, the box spans the interquartile range and the whiskers span the full range, with individual biological replicates overlaid. (+) denotes inflammatory stimulation and (−) its absence.

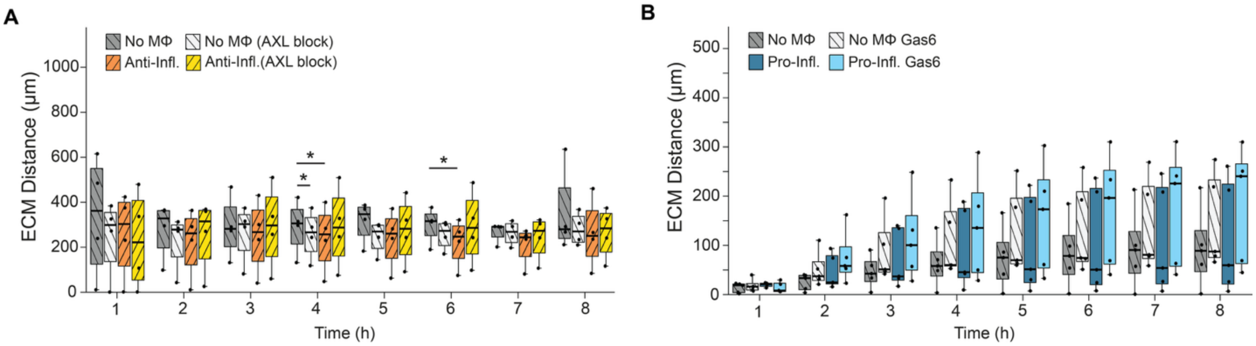


**Fig. S6. Neutrophil migration distance for the Gas6/AXL perturbation experiments.** (A) AXL inhibition, neutrophil migration distance from the lumen edge (µm) over the 8-h window for macrophage-free and anti-inflammatory vessels, each with (AXL block) or without bemcentinib. Asterisks mark the macrophage-free versus anti-inflammatory comparisons at hours 4 and 6, and no comparison between inhibitor-treated and untreated groups reached significance. *$P < 0.05$. $N = 5$ paired, $n = 2$–3. (B) Gas6 addition, neutrophil migration distance from the lumen edge for macrophage-free and pro-inflammatory devices, each with or without exogenous recombinant Gas6 (100 ng/mL). No comparison reached significance at any timepoint ($P > 0.05$). $N = 5$ paired, $n = 2$–3. Paired t-test at each timepoint. In (A) and (B), the center line is the median, the box spans the interquartile range and the whiskers span the full range, with individual biological replicates overlaid. All devices received inflammatory stimulation.
